# A near-complete lamprey genome illuminates ancestral vertebrate innovations

**DOI:** 10.64898/2025.12.15.694513

**Authors:** Zeyu Du, Haixu Wu, Yaoxi He, Jun Li, Qingwei Li, Lin Lin, Jiali Lu, Chunlian Zhu, Kai Liu, Yue Pang, Bing Su

**Affiliations:** College of Life Science, Liaoning Normal University, Dalian, 116081, China; National Key Laboratory of Genetic Evolution and Animal Model, Kunming Institute of Zoology, Chinese Academy of Sciences, Kunming 650201, China; Lamprey Research Center, Liaoning Normal University, Dalian, 116081, China; College of Life Sciences, University of Chinese Academy of Sciences, Beijing 100049, China

**Author notes:** These authors contributed equally (Zeyu Du, Haixu Wu, Yaoxi He).

**Keywords:** Lamprey, *Lethenteron reissneri*, telomere-to-telomere genome, vertebrate origin and evolution, phenotypic innovation, gene family expansion, *CDH2*, heart development, heart chamber formation

## Abstract

The origin of vertebrates is characterized by a suite of phenotypic innovations, yet the underlying genetic mechanisms remain poorly understood. As a living representative of jawless vertebrates, the lamprey serves as a pivotal model for investigating the genomic basis of early vertebrate evolutioin. Here, we report a near telomere-to-telomere (T2T) genome assembly of the reissner lamprey (*Lethenteron reissneri*), generated from sperm DNA that is not subject to programmed genome rearrangement. The 1.25 Gb assembly resolves over 97% of chromosome sequences into single contigs, enabling reconstruction of previously inaccessible genomic regions. Centromeres display highly diversified, lamprey-specific repeat units and structural architectures, whereas telomeres contain both conserved and lineage-specific satellite repeats. Deep transcriptomic profiling across representative organs and developmental stages substantially improved genome annotation. Comparative genomic analyses revealed a major expansion of 1,562 gene families at the base of the vertebrate lineage, and they are significantly enriched for functions related to neural development, skeletal formation, and cardiac function, echoing the known phenotypic innovations in vertebrates. Notably, the *CDH2* gene family underwent vertebrate-specific copy number expansion and acquired conserved cis-regulatory elements (CREs), highlighting its potential role in shaping the early vertebrate circulatory system. Histological analyses and functional knockout experiments demonstrated that one *CDH2* paralog (*CDH2_H*) is essential for lamprey heart development and chamber formation, playing a critical role in the emergence of the closed circulatory system in early vertebrates. This high-quality lamprey genome provides a foundational resource for dissecting the genetic basis of key innovations that shaped vertebrate evolution.

## INTRODUCTION

The emergence of vertebrates approximately 500 million years ago marked a pivotal milestone in the history of life, coinciding with key innovations such as a complex brain, a vertebral skeleton, and a closed circulatory system (1–4). These evolutionary advancements were reportedly driven by two rounds of whole-genome duplication (WGD), which introduced genomic complexity that provided the raw material for vertebrate diversification and rapid radiation (5).

As representatives of living jawless vertebrates (cyclostome), lampreys occupy a critical phylogenetic position for studying early vertebrate evolution (6, 7). For example, lampreys possess a simplified form of two-chambered hearts, likely representing a prototype for the closed circulatory system in the last common ancestor of vertebrates (8). To dissect the genetic basis of such vertebrate innovations, a high-quality reference genome is essential. Although the sea lamprey (*Petromyzon marinus*) genome has been extensively studied (9–11), the current reference remains incomplete, largely due to programmed genome rearrangement (PGR) during development, in which up to 20% of the genome is selectively eliminated from somatic cells (12–16). Thus, only germ cells, such as sperm, retain the complete genome (17). Additionally, the high content of repetitive elements and GC-rich microchromosomes in lampreys presents substantial challenges for producing contiguous genome assemblies (18–21). Fortunately, recent advances in long-read sequencing technologies have provided powerful tools that have been successfully applied to humans and laboratory animals to generate telomere-to-telomere (T2T) genome assemblies (22–29).

In this study, leveraging state-of-the-art long-read sequencing technologies, we generated a high-resolution germline genome assembly of the reissner lamprey (*Lethenteron reissneri*), achieving substantial improvements in completeness and continuity. Furthermore, based on comparative genomic analyses, we reconstructed the events of gene family expansions in vertebrate lineages and uncovered key genes implicated in the emergence of the closed circulating system. These findings deepen our understanding of the molecular mechanisms that shaped vertebrate complexity and adaptive evolution.

## RESULTS

### De novo assembly of the telomere-to-telomere germline genome of the reissner lamprey

Lamprey undergoes genome-wide structural rearrangement during early embryogenesis, resulting in the deletion of approximately 20% (∼0.25 Gb) of the germline genome from somatic lineages. We first tested the genome size of germline and multiple somatic tissues by flow cytometry. As expected, the genome integrity of sperm (∼1.20 Gb genome) is the highest, compared to testis (∼1.08 Gb genome) and other seven types of somatic tissues (∼0.95 Gb genome) (Figure 1A). This suggests that even the testes tissue still contain a large portion of somatic cells subject to PGR. To overcome this challenge, we extracted genomic DNA from sperm cells of a single adult lamprey for genome sequencing (Figure 1B, figure S1, figure S2A). We generated 91.4× Pacific Biosciences (PacBio) high-fidelity (HiFi) reads, 111.4× ultra-long Oxford Nanopore Technology (ONT) reads (≥100Kb), 101.9× high-throughput chromatin conformation capture (Hi-C) reads, and 100.6× Illumina paired-end reads (Figure 1B, figure S1, figure S2B, table S1). For genome annotation, we also produced 600 Gb of short-read RNA-seq and 1,100 Gb of long-read Iso-Seq data across eight tissues and four developmental stages, (Figure 1B, figure S1, table S2; data S1; Methods), enabling comprehensive transcriptome coverage and high-quality genome annotation.

**Figure 1.**
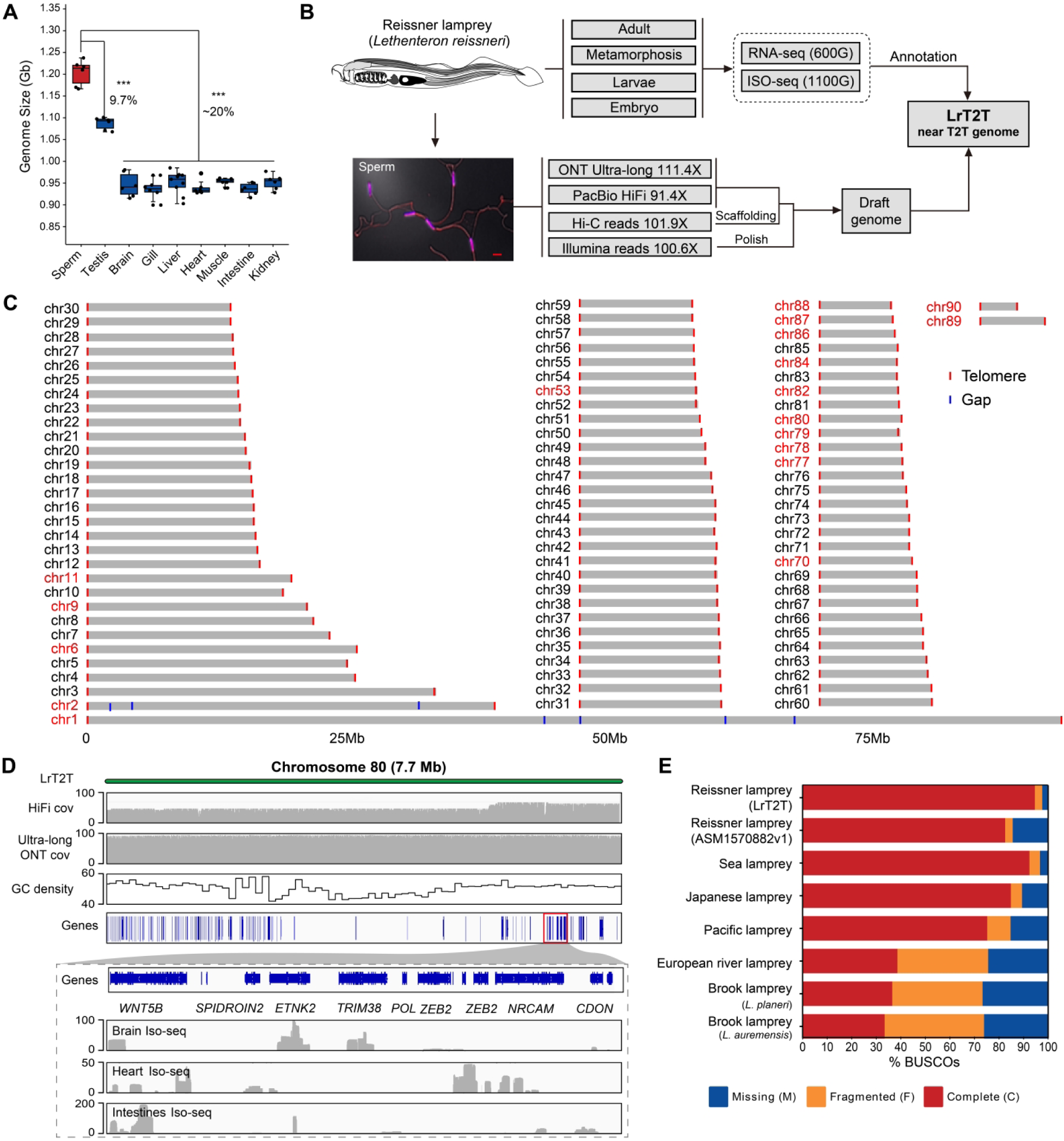
Sequencing and de novo assembly of the germline genome of the reissner lamprey. (**A**) Genome size estimation of germline (sperm and testis) and somatic tissues based on flow cytometry analysis. (**B**) Data-generating pipeline for constructing a near telomere-to-telomere (T2T) genome, integrating long-read sequencing with deep RNA-seq for transcript-supported annotation. ONT Ultra-long, Oxford Nanopore Technology ultra-long ONT reads (≥100Kb). (**C**) Chromosomal distribution of telomeric sequences (red vertical line), and the remaining seven unresolved gaps (blue vertical line) on chr1 and chr2. A total of 18 chromosomes are newly assembled to completion and they are highlighted in red. (**D**) Detailed view of the complete assembly of chromosome 80, validated by high-depth PacBio HiFi and ONT reads, with GC content, gene density, and gene expression (Iso-seq) tracks. cov, coverage. (**E**) Comparison of genome assembly completeness based on BUSCO analysis between LrT2T and previously published lamprey genomes, demonstrating a substantial improvement.

Using hifiasm (v0.20.0-r639) and Verkko (v2.2.1), we assembled a near telomere-to-telomere (T2T) genome (named as LrT2T) of the reissner lamprey. A flow chart for genome assembly and annotation, including the tools used is presented in figure S1. Briefly, with hifiasm, we assembled one main diploid genome, and two haplotype-resolved genomes (table S3). The main assembly showing the highest genome integrity was used for subsequent analysis. In parallel, contigs generated by Verkko were used to resolve residual gaps and improve overall contiguity (figure S1, table S3). The resulting genome spans 1.25 Gb (Table 1), consistent with nuclear DNA content estimated by flow cytometry from post-meiotic sperms (Figure 1A). The contig N50 of the LrT2T assembly is 12.6 Mb with a high quality value (QV) of 54.5 (Table 1, table S4). Based on Hi-C data and manual curation, we scaffolded all contigs into 90 chromosomes (Figure 1C, figure S2C, figure S3, table S5). Notably, over 97 % of chromosomes (88 out of 90) are composed of single contigs without any gaps, indicating exceptional assembly continuity (Figure 1C). Compared to the previous long-read-based genome assembly of reissner lamprey (ASM1570882v1) (13), the LrT2T assembly resolved 18 additional complete chromosomes, many of which span complex regions with high GC content or dense transposable elements (TEs) (figure S3, Table 1, table S6). For instance, although chromosome 80 exhibits the lowest gene density, Iso-seq data confirm that genes on this chromosome are transcriptionally active in the brain, heart, and intestines (Figure 1D), underscoring the accuracy of the assembly and the presence of functional genes within these previously unresolved genomic regions.

**Table 1.**
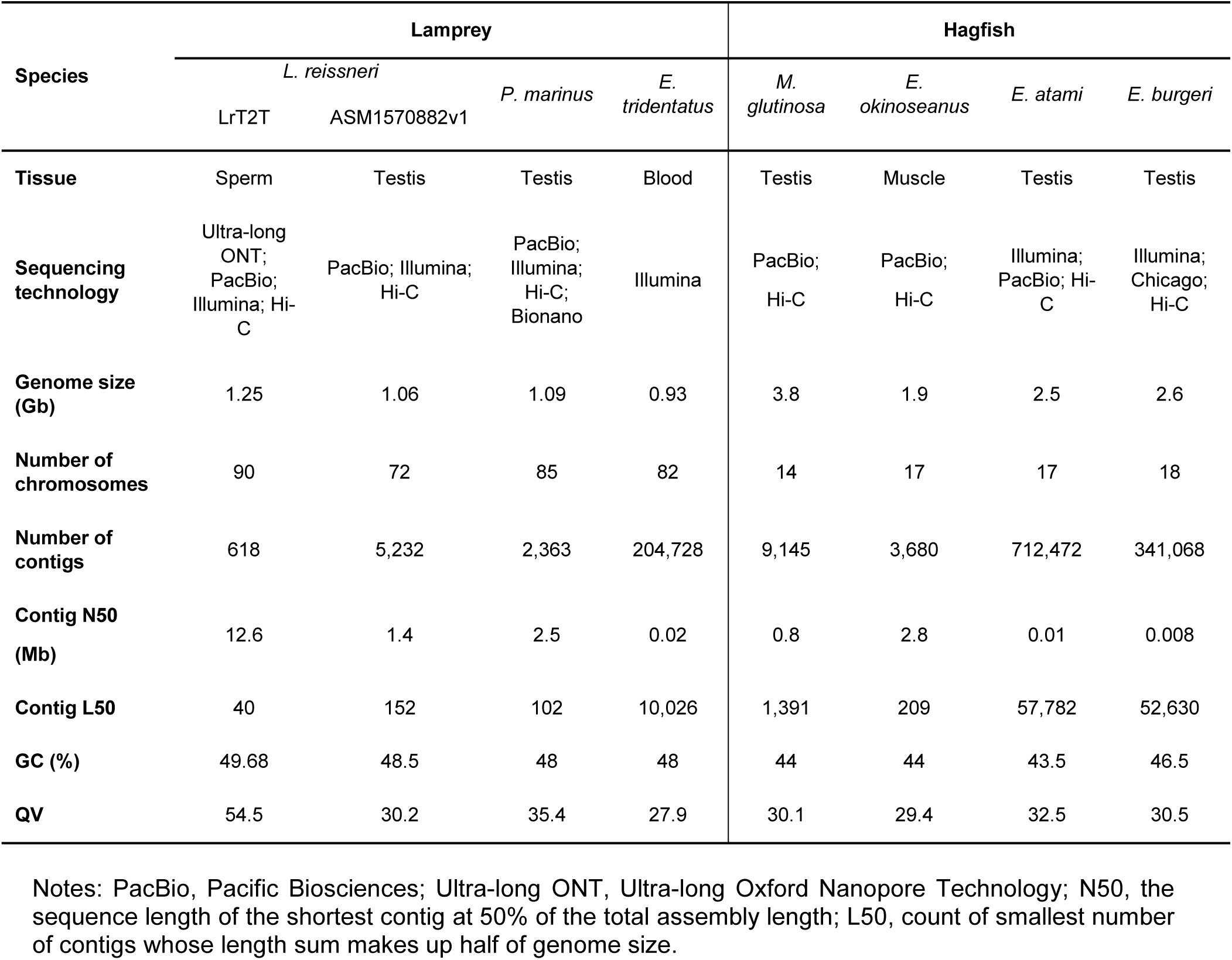
Comparison of LrT2T with the released genomes of jawless fishes.

| Species | Lamprey |  |  |  | Hagfish |  |  |  |
| --- | --- | --- | --- | --- | --- | --- | --- | --- |
|  | *L. reissneri* |  | *P. marinus* | *E. tridentatus* | *M. glutinosa* | *E. okinozeanus* | *E. atami* | *E. burgeri* |
|  | LrT2T | ASM1570882v1 |  |  |  |  |  |  |
| Tissue | Sperm | Testis | Testis | Blood | Testis | Muscle | Testis | Testis |
| Sequencing technology | Ultra-long ONT; PacBio; Illumina; Hi-C | PacBio; Illumina; Hi-C | PacBio; Illumina; Hi-C; Bionano | Illumina | PacBio; Hi-C | PacBio; Hi-C | Illumina; PacBio; Hi-C | Illumina; Chicago; Hi-C |
| Genome size (Gb) | 1.25 | 1.06 | 1.09 | 0.93 | 3.8 | 1.9 | 2.5 | 2.6 |
| Number of chromosomes | 90 | 72 | 85 | 82 | 14 | 17 | 17 | 18 |
| Number of contigs | 618 | 5,232 | 2,363 | 204,728 | 9,145 | 3,680 | 712,472 | 341,068 |
| Contig N50 (Mb) | 12.6 | 1.4 | 2.5 | 0.02 | 0.8 | 2.8 | 0.01 | 0.008 |
| Contig L50 | 40 | 152 | 102 | 10,026 | 1,391 | 209 | 57,782 | 52,630 |
| GC (%) | 49.68 | 48.5 | 48 | 48 | 44 | 44 | 43.5 | 46.5 |
| QV | 54.5 | 30.2 | 35.4 | 27.9 | 30.1 | 29.4 | 32.5 | 30.5 |

Notes: PacBio, Pacific Biosciences; Ultra-long ONT, Ultra-long Oxford Nanopore Technology; N50, the sequence length of the shortest contig at 50% of the total assembly length; L50, count of smallest number of contigs whose length sum makes up half of genome size.

In particular, the high-depth ONT reads helped closing gaps in the previous genome assembly, enabled the annotation of additional genes, and corrected chromosome assembly errors (figure S4). For example, LrT2T chromosome 30 spans a region previously uncovered by HiFi reads, and genes located within this gap exhibit consistently high expression across multiple tissues (figure S4A-B). Additionally, we found that LrT2T chromosome 3 merges two separate chromosomes from the previous assembly (figure S4C). Analysis of microsatellite sequences revealed that the centromeric region was not detected in the earlier version, resulting in misassembly.

Collectively, the LrT2T genome represents a major improvement in completeness, continuity and base accuracy relative to previously published genomes of jawless vertebrates, including both lampreys and hagfishes (13, 16, 30) (Figure 1E, Table 1, table S7).

### Annotation of the LrT2T genome

To facilitate the use of LrT2T in genetic and functional studies, we generated both short-read and long-read RNA sequencing data for genome annotation (Methods), encompassing eight tissue types and four key developmental stages (Figure 1B, table S2). In total, 25,049 genes were annotated. Notably, 99.41% of annotated genes contain complete coding DNA sequences (CDS), compared to only 80.80% in the previous ASM1570882v1 assembly, indicating a substantial improvement in quality and utility of the LrT2T genome (Figure 2A). Importantly, we resolved approximately 190 Mb of previously unresolved regions in the ASM1570882v1 assembly, representing 15.8 % of the LrT2T assembly, including centromeric satellites, various repetitive sequences, and coding sequences (Figure 2B).

**Figure 2.**
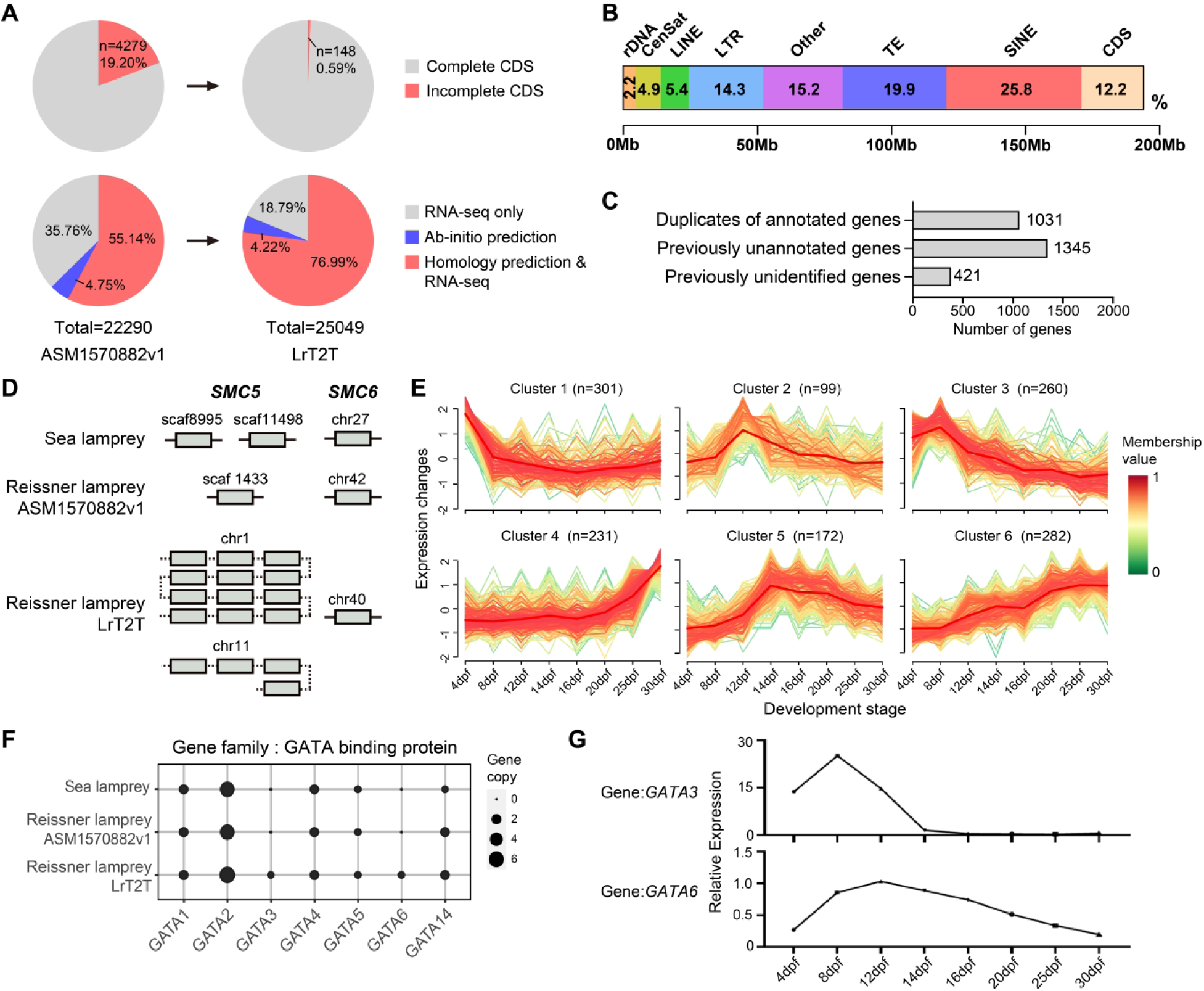
Integrity evaluation of LrT2T genome annotation. (**A**) Comparison of annotated gene numbers, gene integrity (percentage of genes with complete coding sequences, CDS), and supporting annotation evidence between the previous assembly (ASM1570882v1) and the LrT2T genome of the reissner lamprey. Complete CDS indicates that gene sequence is continuous and complete from the start codon to the stop codon. (**B**) Annotations of ∼190 Mb previously unresolved genomic regions, including ribosomal DNA (rDNA), centromeric satellites (CenSat), various repetitive sequences (LTRs, TEs and SINEs), and coding sequence (CDS). (**C**) Classification of the 2,797 additional annotated genes in the LrT2T genome compared to the sea lamprey genome. Among them, “unannotated” refers to genes in the database that have no defined names or functions (only having numbers), while “unidentified” refers to genes that were not covered in the previous genome assembly due to gaps. (**D**) The newly annotated *SMC5* duplicates in the LrT2T genome, located on chromosome 1 (12 copies) and chromosome 11 (4 copies), both newly assembled chromosomes. *SMC6* remains a single-copy gene across all three genome assemblies. (**E**) Expression trajectories of 1,345 previously unannotated genes during embryonic development, grouped into six expression clusters, spanning from 4 to 30 days post-fertilization (dpf). n, gene counts. (**F**) Annotation of *GATA* family genes, among which *GATA3* and *GATA6* are newly identified in the LrT2T genome. (**G**) Expression dynamics of *GATA3* and *GATA6* across embryonic development in the reissner lamprey, spanning from 4 to 30 dpf.

Moreover, compared with the widely used sea lamprey genome (GCF_010993605.1), LrT2T contains 2,797 additional annotated genes (Figure 2C), including duplicated genes previously unannotated (1,031 genes), previously unannotated genes (1,345 genes) and previously unidentified genes due to assembly gaps (421 genes). Notably, a large portion of these additional annotated genes are located on chromosomes 1, 2, 6, and 11. For instance, the SMC5 and SMC6 proteins form a heteromeric complex critical for maintaining chromosomal stability (31). While *SMC6* is consistently present as a single-copy gene across all three lamprey genome assemblies, the copy number of *SMC5* varies — only one copy was annotated in the ASM1570882v1 assembly, and two copies in the sea lamprey genome (Figure 2D). In contrast, LrT2T reveals 16 annotated copies of *SMC5*, with 12 on chromosome 1 and 4 on chromosome 11, both newly assembled chromosomes. Our deep NGS sequencing data (30×) across multiple somatic tissues revealed that 10 *SMC5* copies on Chr1 were absent in these somatic tissues, and they are likely germline-specific, suggesting their potential role in genome stability and programmed genome rearrangement (figure S5A).

Given that the previously unannotated genes (n =1,345) lack functional annotations in current databases, we validated their expression using RNA sequencing data (FPKM >0). We then applied the *mfuzz* tool (32) to analyze their expression dynamics across developmental stages (Methods), and they displayed stage-specific expression trajectories during early embryonic development and post-hatching organogenesis (Figure 2E), suggesting potential roles in developmental processes. Expression of previously unidentified genes (n = 421) was also confirmed, including functionally important genes. For instance, we newly annotated *GATA3* and *GATA6* (absent in previous lamprey genome assemblies), both cardiac transcription factors of the *GATA* family, which exhibited stage-specific expression trajectories during development (from 4 to 30 days post-fertilization, dpf) in the reissner lamprey (Figure 2F-G).

Of note, *GATA4/5/6* exist as a single gene in the sea squirt genome (33). Functionally specialized *GATA* factors appear to have diversified after the divergence of lancelet, potentially contributing to cardiac development. Consistently, our phylogenetic analysis suggests that *GATA4* underwent cyclostome-specific gene duplication (figure S5B). Previous studies in chicken and human have shown that *GATA4* regulates cardiac morphogenesis by transactivation of the *CDH2* gene, and *GATA6* expression was positively associated with *CDH2* expression (34, 35). Therefore, both the extensive expansion of *GATA4* in lampreys and the newly annotated *GATA6* copy may play critical roles in heart formation.

### Assembly and characterization of centromeres and telomeres in the LrT2T genome

Relatively little is known about centromere and telomere organization in non-model animals, largely because these regions consist of extensive arrays of repetitive sequences that are difficult to resolve using short-read sequencing data. In this study, the combination of PacBio HiFi and ONT ultra-long reads enabled the assembly of a single large scaffold for nearly every chromosome, spanning from telomere to telomere (Figure 1C). This high-contiguity assembly allowed us to reconstruct centromeric and telomeric regions across all chromosomes in the LrT2T genome (Figure 3A, Methods).

**Figure 3.**
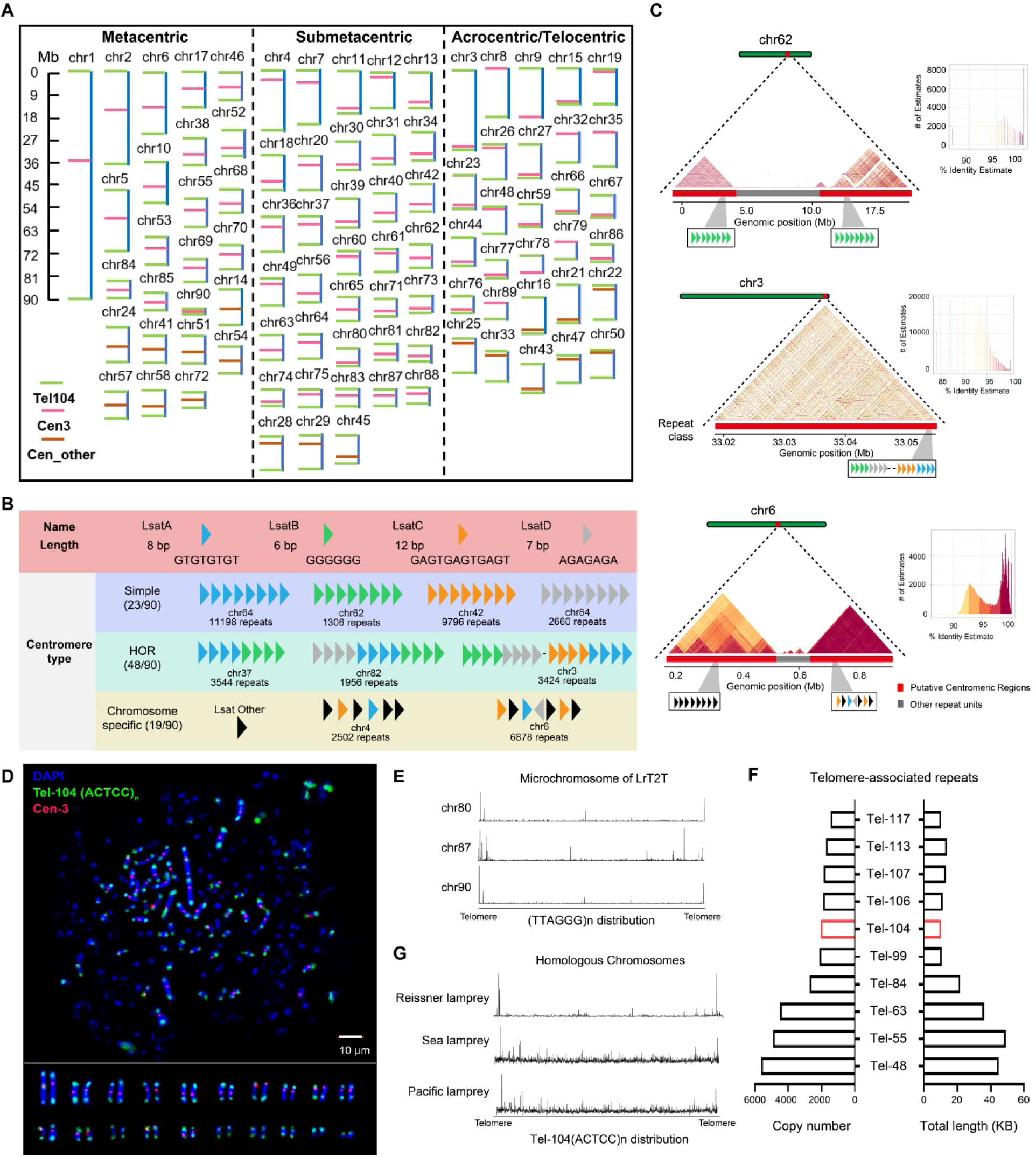
Assembly and characterization of centromeric and telomeric regions in the LrT2T genome. (**A**) The schematic diagrams of 90 chromosomes of the LrT2T genome with resolved centromeres and telomeres. Based on centromere positions, chromosomes are categorized as metacentric, submetacentric, or acrocentric / telocentric. Putative centromeric (Cen-3) and telomere-associated (Tel-104) repeats are highlighted. The scale bar at the top left indicates chromosome size. (**B**) Four repeat families were identified within reissner lamprey centromeres: LsatA (8 bp), LsatB (6 bp), LsatC (12 bp), and LsatD (7 bp). Centromeres contain one to four repeat arrays and can be classified into three architectural categories: simple, HOR (high-order repeat), and chromosome-specific. Simple centromeres contain a single array type, whereas HOR-type centromeres consist of multiple repeat arrays. Repeat units within an array typically share the same orientation; however, in certain chromosomes both orientations coexist, wherein large forward-oriented blocks are interspersed with reverse-oriented repeat units (for example, the LsatD array on chromosome 6). (**C**) Representative centromeric structures from chromosome 62 (simple), chromosome 3 (HOR), and chromosome 6 (chromosome-specific) are shown, together with sequence similarity profiles. Tandem repeat units defining each centromeric region are displayed, and the genomic distribution of Cen-3 and Cen-3-related repeats is illustrated on each chromosome. (**D**) Fluorescent in situ hybridization (FISH) image validating the chromosomal locations of the inferred centromeric repeat Cen-3 and telomere-associated repeat Tel-104 repeats in primary testis cells. Homologous pairs of the 20 relatively large chromosomes are arranged and displayed along the bottom of the panel. (**E**) Distribution of vertebrate-conserved telomeric sequences (TTAGGG)_n_ across three microchromosomes (difficult to assemble chr80, chr87, and chr90) in the LrT2T genome. The height of the peak represents the number of repeats of the motif. (**F**) Total length and copy number of the top 10 most abundant telomere-associated satellite repeats. (**G**) Distribution of the Tel-104 repeat across homologous chromosomes in three lamprey genome assemblies. The height of the peak represents the number of repeats of the motif.

To systematically identify centromeric and telomeric regions, using multiple tools including CentIER (v2.0), TRASH (v1.0), and TIDK (v0.2.1), we quantified sequence repetitiveness using entropy and linguistic complexity metrics, yielding candidate repeat-rich domains across all chromosomes (Methods). For centromere identification, rather than relying solely on tandem repeat detection, we implemented an integrative framework that considers multiple sequence features, including tandem repeat density, retrotransposon content, and k-mer frequency distributions. These highly repetitive domains ranged from less than 1 Mb to over 10 Mb in length. Representative repeats from these regions were identified and selected to define centromeres and telomeres. Complete sequences of the inferred centromeric and telomeric repeats are provided in data S2 and data S3.

For non-model vertebrate species, defining conserved centromeric features is challenging, as no uniform repeat architecture or shared consensus sequence can be clearly identified, despite the presence of tandem repeat arrays on every chromosome. We identified four dominant simple repeat units—LsatA, LsatB, LsatC, and LsatD, which form distinct array types within the inferred centromeric regions (Figure 3B; figure S6). These Lsat elements are arranged into one of four structural arrays that define simple-type centromeres (Figure 3B; figure S6). In contrast, HOR (high-order repeat) centromeres represent the most prevalent architecture, characterized by mixed repeat blocks that interweave to form multi-array structures (Figure 3B; figure S6). Additionally, at least 19 chromosomes possess centromeres that cannot be represented by simple repeat arrangements and instead show strong chromosome-specific signatures, exemplified by the unique organization of chromosome 6 (Figure 3B; figure S6). The HOR centromeres are typically 35 to 235 Kb in length, while the simple and chromosome specific centromeric regions are longer and show higher sequence similarity (Figure 3C, data S2).

We identified a 48-bp repeat unit (named Cen-3) in the centromeric region of chromosome 3 (a telocentric chromosome) (Figure 3B-C). Homologous search detected Cen-3-related repeat units (LsatA/B/C/D) in the centromeric regions of most chromosomes (71 out of 89 chromosomes), and different chromosomes exhibit length and sequence variations. For example, in chromosome 42, the dominant portion consists of GAGTGAGTGAGT repeat regions (LsatC), whereas in chromosome 62, the repeats contain a higher proportion of G (LsatB) (Figure 3B). These Cen-3-related repeats represent a centromere organization markedly distinct from the typical alpha-satellite DNA observed in jawed vertebrates. In contrast, the centromeric repeat units in other chromosomes that do not have Cen-3-related repeats are highly variable. To evaluate whether the predicted repetitive arrays correspond to active centromeric regions, we performed immunodetection assays using commercially available mouse-derived CENPA (centromeric protein A) antibodies. However, the antibody failed to recognize centromeres in lamprey cells (figure S7), preventing direct validation at the protein level. In contrast, our fluorescence in situ hybridization (FISH) experiments confirmed the genomic locations of the centromeric repeat Cen-3 on multiple chromosomes (Figure 3D), supporting the reliability of the identified centromeric regions and their repeat units. Together, these results indicate that in lamprey, centromeric regions and their sequence features exhibit extensive variability (17).

For telomeres, lampreys exhibit conserved sequence features shared with jawed vertebrates, namely telomeric satellite repeats composed of (TTAGGG)_n_, and all 90 assembled chromosomes including those small-sized micro-chromosomes contain telomeres at both ends (Figure 3E, figure S8). In addition, the telomeric regions in lampreys, defined as the terminal 1% at both ends of each chromosome, contain a greater diversity and higher abundance of 4-8 bp tandem repeat arrays. The sequence complexity and repeat frequency in these regions are markedly higher than those observed in jawed vertebrates (Figure 3F, data S3). This divergence may be associated with evolutionary events involving chromosomal breakage and the formation of micro-chromosomes in lampreys. For instance, the telomere-associated satellite repeat Tel-104 is lamprey-specific and exhibits high sequence specificity, with strict localization at both chromosomal ends (Figure 3G). Our FISH experiments confirmed the genomic positions of the telomere-associated repeat Tel-104, demonstrating their presence in telomere structure (Figure 3D).

### Identification of gene family expansions and core genes driving the evolution of the vertebrate circulatory system

Four whole-genome duplication (WGD) events are inferred to have occurred during vertebrate evolution: in the vertebrate common ancestor (1R), in jawed vertebrates (2R), in teleost fish (3R), and in the common ancestor of living jawless vertebrates (CR) (36). These WGD events are believed to have had a profound impact on vertebrate phenotypic innovations (37). Using the LrT2T genome, we performed a comparative genomic analysis to identify lineage-specific gene family expansions (Methods), incorporating high-quality genome sequence data from 75 representative vertebrate species. The genomes of lancelet and sea squirt were also included as outgroups (Methods) (15, 20, 27, 28, 38).

We detected extensive gene family expansions across the vertebrate phylogeny, particularly in lineages that experienced WGD events (Figure 4A). In the vertebrate common ancestor, we identified 1,562 lineage-specific gene families (Figure 4A, data S4, methods), likely resulting from the 1R event. Of note, the newly annotated genes in LrT2T include a substantial number of these vertebrate-specific duplicates, accounting for about 30% of the identified 1,562 vertebrate-specific gene families, indicating the necessity of generating a complete reference genome. We thus focused on these 1,562 gene families to evaluate their potential contribution to phenotypic innovations at the origin of vertebrates. Gene Ontology (GO) enrichment analysis revealed that these gene families are significantly enriched for functions related to neural development (figure S9A), skeletal formation (figure S9B-C), and cardiac function (Figure 4B, data S5), echoing the known phenotypic innovations in vertebrates. The most enriched is cardiac function, including intercalated disc, fascia adherent, cell-cell junction (figure S10A) and other heart-related functions (figure S10B). Given that differentiation of atrial and ventricular structures represents a key evolutionary innovation in vertebrates leading to the emergence of the closed circulating system (multichambered heart) in vertebrates, the enrichment of cardiac-related functions among vertebrate-specific gene families suggests that newly evolved genes may have contributed to this innovation.

**Figure 4.**
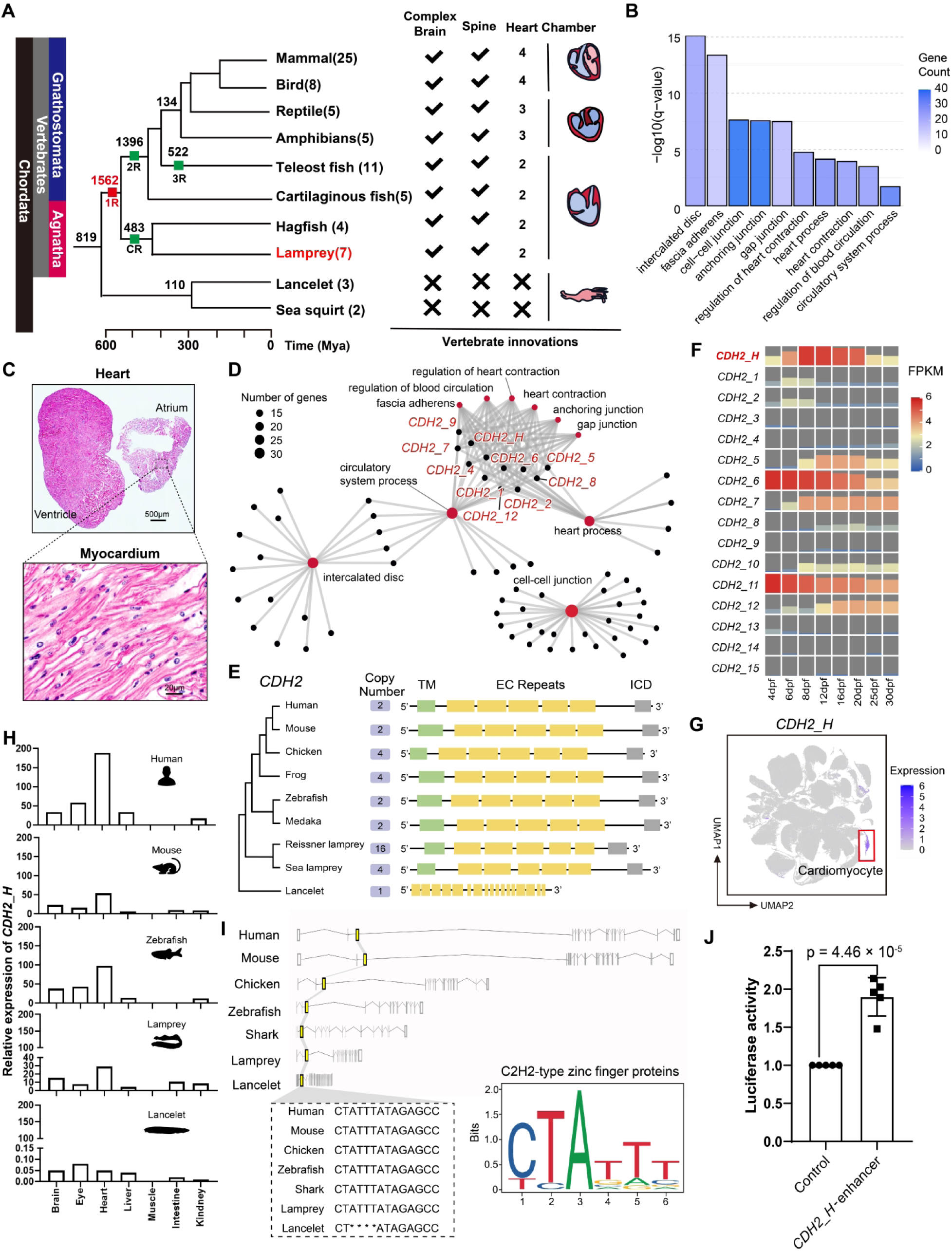
Identification of lineage-specific gene family expansions in vertebrates and characterization of the vertebrate-specific *CDH2* expansion. (**A**) Lineage-specific gene family expansions mapped onto the vertebrate phylogeny. Major expansions coincide with proposed whole-genome duplication (WGD) events, including those in the last common ancestor of vertebrates (1R), jawed vertebrates (2R), teleost fishes (3R), and the common ancestor of extant jawless vertebrates (CR). Lancelet and sea squirt species serve as outgroups. The numbers in parentheses following the major vertebrate lineages indicate the number of species and their genomes collected from the published data. A total of 75 genomes were used for the comparative genomic analysis of the vertebrate lineage. The expansion of 1,562 gene families in the vertebrate ancestor is highlighted in red. Key phenotypic innovations during vertebrate evolution are annotated along the phylogeny. An evolutionary timeline is shown at the bottom. (**B**) Significantly enriched Gene Ontology (GO) terms related to cardiac function. (**C**) Hematoxylin and eosin (H&E) staining images showing the two-chambered heart of the lamprey. (**D**) Gene-Concept network derived from 69 gene families enriched in cardiac-related GO terms. The network contains 10 *CDH2* gene copies and they are highlighted in red. (**E**) Evolutionary expansion of the *CDH2* gene family in lampreys and jawed vertebrates (≥2 copies), in contrast to the single copy in lancelets. The conserved functional domains of *CDH2* in vertebrates are illustrated, including transmembrane domain (TM), extracellular cadherin (EC) repeats, and intracellular signaling domains (ICD). (**F**) Developmental expression patterns of the 16 *CDH2* gene copies in lamprey embryos across 8 developmental stages (4, 6, 8, 12, 16, 20, 25, and 30 dpf). *CDH2_H* shows elevated expression during the critical window of heart development (12 dpf). (**G**) Single-cell transcriptomic analysis indicates that *CDH2_H* is specifically expressed in cardiomyocytes. (**H**) Conserved heart-specific expression of *CDH2_H* in vertebrates including lamprey, zebrafish, mouse, and human. The expression level in lancelet is much lower. (**I**) A vertebrate-specific enhancer located within the intronic region of *CDH2_H*, containing a highly conserved motif predicted to be bound by C2H2-type zinc finger proteins. (**J**) Functional validation of the *CDH2_H* enhancer by luciferase reporter assays in primary cultured mouse cardiomyocytes (PCMCM). Data are shown as mean ± SEM from at least five independent experiments. Statistical significance was assessed by two-way ANOVA with Tukey’s multiple comparison test.

Using hematoxylin and eosin (H&E) staining, we confirmed that the reissner lamprey indeed possesses a two-chambered heart (Figure 4C). To explore the functional modules likely involved in emergence of multichambered heart, we generated Gene-Concept networks of 69 genes with vertebrate-specific expansion and are enriched in cardiac-related GO terms (Figure 4B, Methods). The network pattern suggest that many genes are functionally connected (Figure 4D). These include *SLC8A3*, *SLC25A23*, and *HCN2*, which play key roles in Ca²⁺ transport and heart physiology (39–41).

Notably, there are ten *CDH2* gene copies in the network (Figure 4D, data S6). Comparative genomic analysis revealed at least two *CDH2* copies in all analyzed vertebrate species, whereas in invertebrates such as lancelets, only a single copy is present, suggesting that WGD events may have driven *CDH2* expansion during vertebrate evolution (Figure 4E, figure S9D). Although *CDH2* was also detected in the sea lamprey genome assembly, only four copies were identified, in contrast to the sixteen *CDH2* copies seen in the LrT2T assembly (42). These 16 copies are scattered across different chromosomes, likely resulting from the vertebrate-shared 1R, the lamprey-hagfish-shared CR, and gene family duplication events (figure S11, figure S12), although the linkage patterns of these *CDH2* copies do not show clear sign of segmental duplications (figure S11). The sequence similarity among the 16 *CDH2* copies of *L. reissneri* ranges from 33.98%-98.50%. Taking the lancelet *CDH2* copy as outgroup, we constructed a gene family tree of *CDH2* copies, which places *CDH2_H* near the root of the gene family (figure S12). Gene structure analysis shows high conservation of *CDH2* functional domains across vertebrates, including five extracellular cadherin repeats, and one intracellular signaling domain, which are also seen in most of the *CDH2* copies of reissner lamprey, while in lancelet, the functional domains are incomplete (Figure 4E, figure S12). Given these findings, we speculate that *CDH2* expansion may have contributed to the emergence of the multichambered heart in the vertebrate ancestor.

Besides of *CDH2*, we also detected pronounced expansion in multiple cardiac function–associated gene families, including *FGF6*, *KCNIP2/4*, *MYH7*, *PTPR2*, and *TLE3/4* (43–47). These families encode proteins involved in cell junction formation, signal transduction in cardiomyocytes, and cardiac developmental regulation, and both gene copy number and protein domain architectures show lineage-dependent evolutionary diversification (figure S10C, figure S13). Furthermore, we found that several canonical transcription factors essential for cardiac chamber formation exhibit distinct evolutionary signatures. Specifically, *TBX5* and *HAND2* demonstrate cyclostomata-specific gene expansion, whereas *NR2F2* and *ZNF711* represent newly evolved genes in vertebrates (figure S14) (48–51). Taken together, these observations indicate that the emergence of cardiac chambers in vertebrates was likely shaped by the combined contributions of vertically inherited genes, 1R/CR-derived gene expansions, and de novo gene formation during vertebrate origin and early diversification.

To evaluate the functional roles of *CDH2* gene copies in lampreys, by sampling the entire embryos, we examined their expression patterns across 8 embryonic developmental stages (4 dpf - 30 dpf). Most *CDH2* copies were expressed during development (Figure 4F). Notably, one copy, designated *CDH2_H*, a known cardiac *CDH2* homolog in jawed vertebrates (52), was highly expressed at 12 dpf (T24-25, hatching and heart beating), a critical stage for heart development in lamprey (53). We further analyzed *CDH2_H* expression using published single-cell RNA-seq data from pre-metamorphic larval lampreys (2.5–3 years old) (54). Among 70 cell types across 14 tissues, *CDH2_H* showed specific and robust expression in cardiomyocytes (Figure 4G), whereas other *CDH2* copies lacked such specificity (figure S15). This heart-specific expression pattern is conserved in both lampreys and jawed vertebrates, including zebrafish, mouse, and human, while its expression in lancelet is extremely low (Figure 4H), further supporting a conserved role for *CDH2_H* in vertebrate heart development.

To investigate the regulatory mechanisms underlying *CDH2_H* expression, we performed comparative genomic analyses to identify conserved cis-regulatory elements (CREs) across vertebrate genomes. We identified a vertebrate-specific enhancer located in an intronic region of *CDH2_H* (Figure 4I). This enhancer harbors a conserved binding motif for C2H2-type zinc finger proteins, transcription factors implicated in cardiac development and congenital heart disease (55). This motif is conserved in vertebrates but absent in lancelet. To functionally validate the enhancer, we conducted dual-luciferase reporter assays in primary cultured mouse cardiomyocytes (PCMCM), which confirmed its regulatory activity (Figure 4J). Together, our findings suggest that *CDH2_H*, a vertebrate-specific *CDH2* paralog, may have acquired heart-specific expression through the recruitment of a conserved enhancer. This regulatory innovation likely contributed to the evolution and function of the multichambered heart in early vertebrates.

### Functional disruption of CDH2_H impairs lamprey cardiac development

To investigate the functional role of *CDH2_H* in lamprey heart development, we generated two knockout lines: a *CDH2_H* gene knockout (*CDH2_H*-ko) and an enhancer knockout (enhancer-ko). For each line, three sgRNAs were designed to target either exon 1 (*CDH2_H*-ko) or the intronic enhancer (enhancer-ko) of *CDH2_H* at one-cell stage (Figure 5A, table S8, Methods). The Illumina genome sequencing data (50×) revealed high knockout efficiencies: 94.12 % (48/51) for *CDH2_H*-ko, 89.80 % (44/49) for enhancer-ko—with no detected off-target effect (data S7, data S8). At 6 dpf (T19), no obvious morphological abnormalities or differences in survival rates were observed among the control and knockout embryos (Figure 5B). However, by 12 dpf (T24-25), a key window for cardiac morphogenesis, both mutants displayed markedly increased embryonic lethality, and only 2.4% (*CDH2_H*-ko) and 5.4% (enhancer-ko) of the knockout lampreys survived to 20dpf (T27-28) (Figure 5B). Consistently, RT–qPCR of heart tissues confirmed significant downregulation of *CDH2_H* in both knockouts at 12 dpf (Figure 5C).

**Figure 5.**
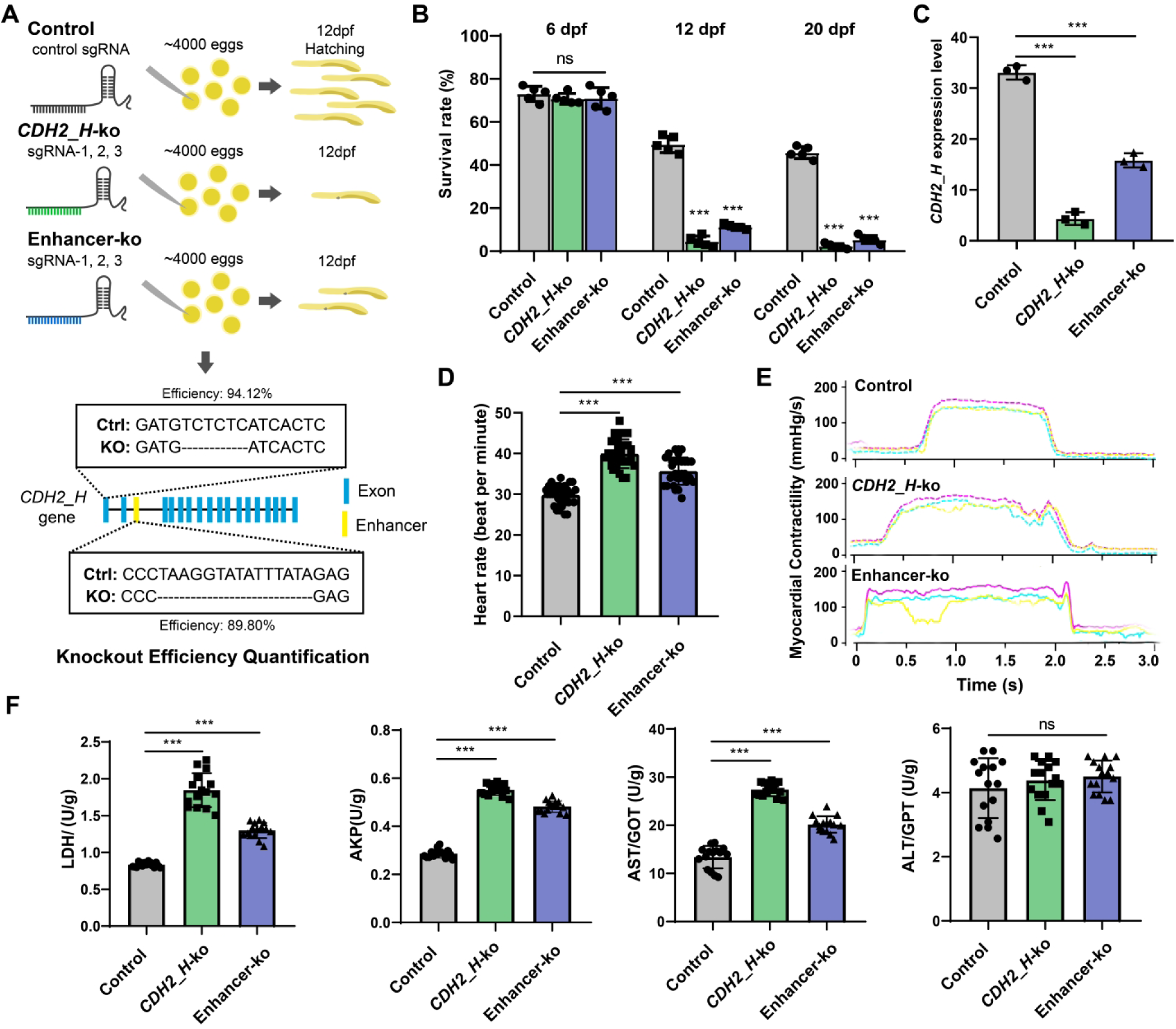
Targeted knockout of *CDH2*_*H* or its enhancer disrupts heart development in lampreys. (**A**) Schematic diagram showing the knockout design of lamprey *CDH2_H* exon-1 and the enhancer located in intron-2. For each knockout line, three sgRNAs were designed. For each group, 4,000 fertilized eggs were used for injection, and each group had three independent experiments (12,000 fertilized eggs in total). Knockout efficiency and loci were demonstrated through genome sequencing of embryos. (**B**) Survival rates of embryos from control and the two knockout lines (*CDH2_H*-ko and enhancer-ko) at three developmental stages (6 dpf, 12 dpf and 20 dpf). n = 500 per group, with 5 independent biological replicates (n = 100). (**C**) *CDH2_H* expression levels in the heart at 12 dpf, quantified by real-time quantitative PCR (RT-qPCR), with 3 independent biological replicates (n = 10). (**D**) Heart rate comparison at 12 dpf among control, *CDH2_H*-ko, and enhancer-ko, with 3 independent biological replicates (n = 30). (E) Electrophysiological recordings at 12 dpf showing differences in myocardial contractility among the three groups. There parameters were recorded, including cardiac contractility (pink line), action potentials (yellow line), and spontaneous ectopic activity (blue line), with 3 independent biological replicates (n = 10). (F) Measurements of three enzymes for cardiac injury evaluation (LDH, AKP, and AST), and liver-specific ALT as a control in 12 dpf lamprey hearts, with 3 independent biological replicates (n = 15). Each data point indicates the measurement of a pooled sample of 10 individuals, and each group contains 15 pooled samples. Values are shown as mean ± SE (standard error) from at least three independent experiments. Statistical analysis was performed using two-way ANOVA with Tukey’s multiple comparisons test. ns, not significant; ***p < 0.001.

Functional assessment of the survived knockouts (12 dpf) revealed clear changes of cardiac functions. Resting heart rates were significantly elevated in the knockouts (39.93 ± 3.46 bpm in *CDH2_H*-ko, 35.67 ± 3.30 bpm in enhancer-ko) compared with controls (29.8 ± 2.43 bpm) (*p*<0.001, two-way ANOVA) (Figure 5D). Electrophysiological recordings revealed enhanced cardiac contractility, prolonged action potentials, and spontaneous ectopic activity in the knockouts, indicative of a pro-arrhythmic phenotype (Figure 5E). The heart tissue biochemical assays revealed significantly elevated levels of lactate dehydrogenase (LDH), alkaline phosphatase (AKP), and aspartate aminotransferase (AST) in both knockout lines, consistent with cardiac damage (Figure 5F). In contrast, alanine aminotransferase (ALT), a liver-specific enzyme, remained unchanged, excluding general hepatic toxicity (Figure 5F). These results demonstrate that *CDH2_H* is essential for proper cardiac development and electrical stability in lamprey, and its disruption leads to severe arrhythmogenic dysfunction and early developmental lethality.

### CDH2_H controls cardiac chamber morphogenesis and underlies the evolutionary emergence of a closed circulatory system in vertebrates

Given the pronounced cardiac dysfunction and lethality observed in *CDH2_H* deficient embryos, we next sought to uncover the structural and regulatory mechanisms responsible for these phenotypes. Among the surviving embryos (12 dpf), pericardial edema, a hallmark of cardiac malformation, was observed in 85.5 % ± 4.9 % of *CDH2_H*-ko (693/814) and 74.0 % ± 1.5 % of enhancer-ko (707/956) embryos, compared to only 2.3 % ± 1.2 % in controls (7/300) (Figure 6A). The mRNA in situ hybridization images show a clear heart-region-specific expression of *CDH2_H* at 12dpf in control lampreys, while no expression was detected in the two knockout lines (Figure 6B). Histological analyses showed disorganized cardiomyocytes with large intercellular spaces in the heart of 12 dpf knockouts, in contrast to the densely packed cardiomyocytes in controls (Figure 6C). More importantly, cardiac chamber defects were observed, where the controls show clear formation of one atrium and one ventricle, while such heart structure got disrupted in the knockouts (Figure 6C). Due to these developmental defects, nearly 90% of the knockout lampreys died before 12dpf (Figure 5B). Meanwhile, no abnormalities were observed in other organs, indicating that the phenotype is restricted to changes during heart development (figure S16). Furthermore, we also performed H&E staining at 20dpf, and the chamber-disruption phenotype was confirmed (Figure 6D).

**Figure 6.**
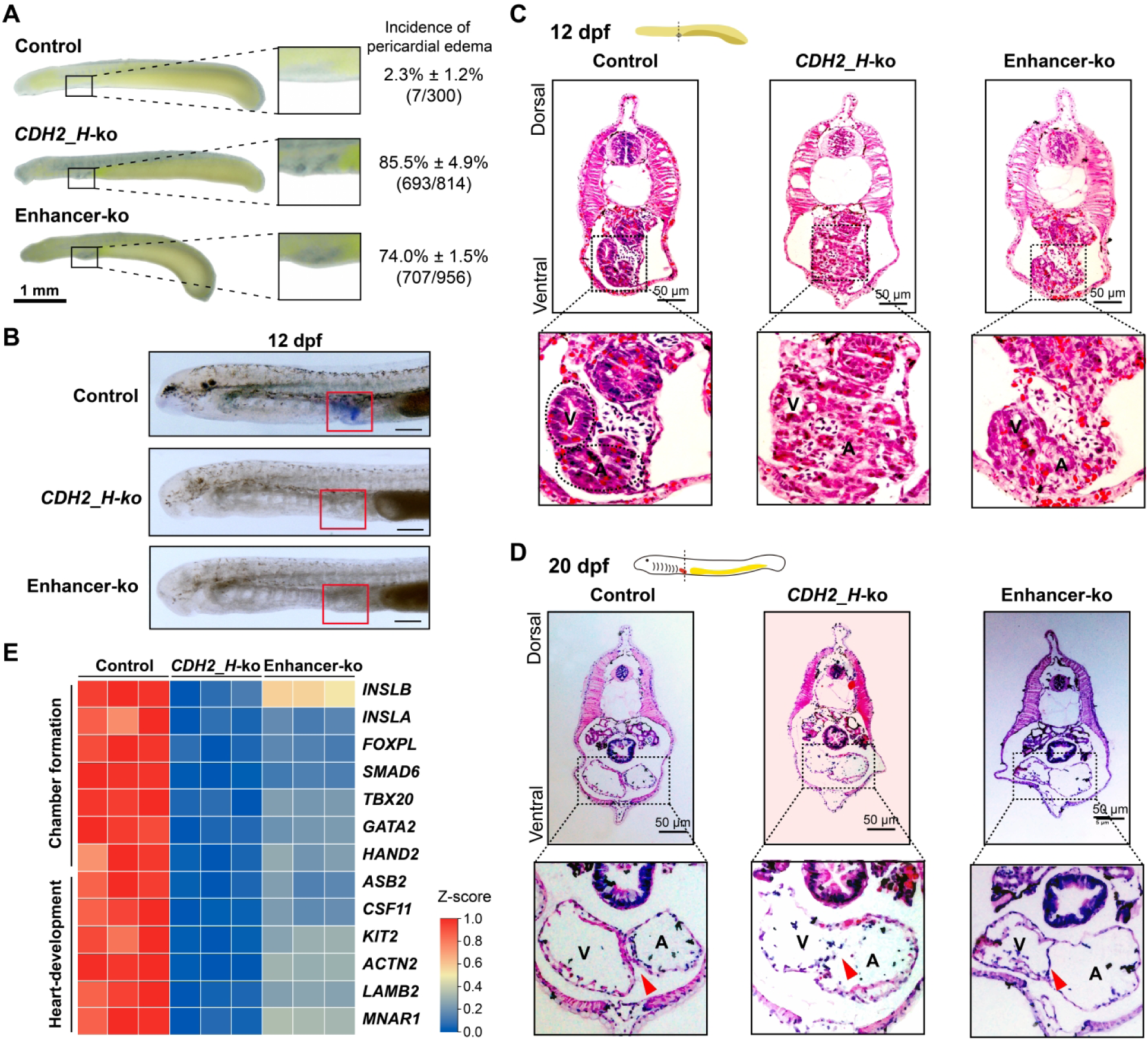
Targeted knockout of *CDH2*_*H* or its enhancer disrupts heart development in lampreys. (**A**) Representative images of 12 dpf embryos showing pericardial edema (a hallmark of abnormal heart development) in the knockout lines. The incidence of pericardial edema was 85.5 % ± 4.9 % in *CDH2_H*-ko (693/814), 74.0 % ± 1.5 % in enhancer-ko (707/956), and 2.3 % ± 1.2 % in controls (7/300). (**B**) Images of mRNA in situ hybridization of *CDH2_H* in 12 dpf lampreys. *CDH2_H* expression was detected in the heart region of control embryos (blue signal within the red-boxed area), whereas no detectable signal was observed in the knockouts. Scale bar, 100 μm. (**C**) Histological assessment of cardiomyocyte organization by hematoxylin and eosin (H&E) staining of 12 dpf hearts. The magnified area is a typical cardiac region, while the atria (A) and ventricles (V) are highlighted, and the scale is 50 μm. (**D**) Histological assessment of cardiomyocyte organization by H&E staining of 20 dpf hearts. The red triangle indicates the septum between the atrium (A) and the ventricle (V). The scale is 50 μm. (**E**) RT-qPCR quantification of 13 genes associated with chamber formation and heart development in 12 dpf lamprey hearts. Relative expression levels are illustrated using color-coded heatmaps. Each group contains three biological replicates.

To test if *CDH2_H* disruption affect expression of other genes, we performed RT-qPCR analysis of 13 known cardiac developmental genes (e.g., *TBX20* and *SMAD6*) (56) at 12 dpf. We observed significant downregulation across all tested genes in the knockout lines (Figure 6E), implying that *CDH2_H* may act as an upstream regulator of heart development in lamprey. Consistently, in mice, conditional knockout of the *CDH2* encoding N-cadherin results in abnormal heart development (57). In humans, heterozygous missense mutation of *CDH2* can cause arrhythmogenic cardiomyopathy/dysplasia (58). Hence, evidence from multiple species support a conserved functional role of *CDH2_H* in lamprey and its homology in jawed vertebrates.

Taken together, these findings indicate that *CDH2_H*, a newly evolved and vertebrate-specific gene copy with a vertebrate-conserved enhancer, plays a pivotal role in cardiac chamber formation and may have contributed to the emergence of the closed circulatory system in the vertebrate common ancestor.

## DISCUSSION

In this study, by leveraging advanced sequencing technologies, we assembled a near-T2T genome of the reissner lamprey, representing a substantial improvement in completeness, continuity and accuracy over currently available genomes of jawless vertebrates. With significantly enhanced gene annotation, the LrT2T genome provides a high-quality reference for the basal clade of vertebrates. Using this assembly, we characterized the organization and sequence diversity of centromeres and telomeres, features not resolved in previously published lamprey genomes. Through comparative genomic analyses across representative vertebrate species, we identified lineage-specific gene family expansions, particularly in the lineage leading to the origin of vertebrates. Among these, we found a vertebrate-specific expansion of the *CDH2* gene family, which we demonstrated to play a critical role in heart development and function, suggesting its involvement in the emergence of a closed circulatory system in the vertebrate common ancestor.

Using the LrT2T genome, we resolved the previously uncertain chromosome number (2n = 144–162) in the reissner lamprey, which had been difficult to determine due to the presence of numerous microchromosomes, challenging to distinguish by traditional cytogenetic methods, and the effects of PGR, which eliminates approximately 20 % of the genome (17, 59). We successfully assembled the genome into 90 chromosomes, each containing both centromeric and telomeric structures, resulting in a diploid genome with 180 chromosomes (2n = 180). This estimate is consistent with the reported chromosome number (2n = 180–198) of the sea lamprey based on long-read genome assembly (60). Notably, despite the near-T2T quality, two chromosomes (chromosomes 1 and 2) still contain several assembly gaps (Figure 1C), likely due to their large size and/or complex sequence content, issues also unresolved in the sea lamprey genome. Together, our results highlight the necessity of using sperm DNA to achieve high-quality T2T genome assemblies for lamprey, given PGR effects in somatic cells and limited availability of oocytes.

In the lamprey genome, the centromeric regions exhibit poorly conserved structural organization and we identified diversified repeat unit sequences. This observation aligns with the notion that, in vertebrates, centromeres are generally composed of tandem repeat arrays, while the length and sequence of the repeat units vary substantially across species (60, 61). Given that the ancestral chordate karyotype is thought to have consisted of 23 large chromosomes (20), we speculate that extensive chromosomal breakage events occurred in the lamprey lineage, giving rise to numerous small, punctate chromosomes. These events likely contributed to the increased structural complexity and sequence variation observed in lamprey centromeres. We attempted to experimentally test the centromere sequence of the reissner lamprey using the CENPA antibody from mice, but unfortunately, the antibody was unable to recognize the centromere region of the lamprey (figure S6). Consequently, generating a lamprey-specific CENPA antibody seems necessary to obtain experimental evidence in future studies. In contrast, lamprey telomeric regions display sequence features conserved with those of jawed vertebrates, although we also identified multiple lamprey-specific telomeric repeat sequences. These repeats may be associated with chromosomal fragmentation and the formation of microchromosomes unique to the lamprey genome, which call for further genomic dissection and experimental validation.

Multiple WGD events occurred during the origin and early evolution of vertebrates, providing a wealth of genetic material that likely contributed to the emergence of vertebrate-defining phenotypic innovations (62). Recent analysis of the hagfish genome, together with phylogenetic data, has offered a more comprehensive view of vertebrate genome evolution (63). By incorporating the newly assembled LrT2T genome, we further clarified the impact of WGD events. More importantly, we identified lineage-specific gene family expansions across key nodes of vertebrate evolution, particularly the expansion of 1,562 gene families in the lineage leading to the last common ancestor of vertebrates. Functional enrichment analyses revealed that these gene families are associated with cardiac function, neural development, and skeletal formation, traits that correspond to hallmark vertebrate innovations such as a closed circulatory system, a complex brain, and a vertebral skeleton. These findings support the hypothesis that gene family expansion played a central role in enabling the emergence of new morphological and physiological features during early vertebrate evolution.

Chordate amphioxus and ascidians possess hearts that consist solely of aortic vessels, whereas lampreys represent the earliest vertebrates with a closed, tubular circulatory system (64). GO enrichment analysis of the expanded gene families in the lamprey genome revealed that many terms associated with heart development are specifically linked to the formation of intercalated discs that connect adjacent cardiomyocytes and are composed of multiple types of junctions, including tight junctions and desmosomes (65). These structures first appeared in lampreys and primarily formed by specialized cell junctions (66).

Our comparative genomic analysis of genes related to intercalated disc connectivity across chordates revealed extensive expansion of this gene superfamily in vertebrates. In contrast to amphioxus and ascidians, vertebrates exhibit clear signs of gene duplication and directional selection in cell junction genes, likely reflecting their functional importance in vertebrate organogenesis. Among the expanded gene families, cell adhesion molecules (CAMs) are particularly notable. These molecules are critical for processes such as cell migration, synapse formation, and target recognition (67). Within this group, cadherins represent a large family of calcium-dependent adhesion proteins that mediate cell-cell cohesion (68). For instance, various cadherins such as *FAT1* and *CDH5* play a role in cell differentiation and are involved in the formation of tissues during embryonic development (69–71). *CDH2*, which encodes N-cadherin, is a type I classical cadherin essential for early brain and heart morphogenesis (72). Studies have shown that the atria and ventricles of zebrafish with *CDH2* mutations are arranged abnormally (51, 73). Meanwhile, in the hearts of *CDH2*-ko mice, the arrangement of muscle fibers at the interventricular septum is disorganized, accompanied by edema and loss of morphology, which leads to the death of the individual (56, 74). Our *CDH2_H* knockout experiments further support its pivotal role in cardiac development and chamber formatioin, suggesting that expansion of the *CDH2* gene family in the vertebrate common ancestor may have contributed to the evolution of a closed circulatory system, namely the multichambered heart.

This study has several limitations. First, although we assembled a near-T2T germline genome for the lamprey, a few assembly gaps remain on two chromosomes (chr1 and chr2). These regions present opportunities for future improvement. Second, our comparative genomic analyses focused primarily on gene family expansions associated with WGD events during early vertebrate evolution. Other important genetic mechanisms, such as programmed genome rearrangement, gene loss, and the role of functional repetitive elements, warrant further investigation. Additionally, while this study primarily focuses on the evolution of the multichambered heart, other key phenotypic innovations during vertebrate origins also merit detailed comparative and functional genomic analyses to fully understand the genetic basis of vertebrate diversification and radiation.

In conclusion, we assembled a high-quality reference genome for a basal vertebrate lineage, with resolved centromeric and telomeric regions. This enabled the identification of lineage-specific gene family expansions, notably that of the *CDH2* gene family, which we demonstrate to be essential for lamprey heart development and function. Our findings suggest that *CDH2* expansion may have contributed to the origin of the multichambered heart in vertebrates. The LrT2T genome represents a valuable resource for advancing our understanding of vertebrate evolution and the genetic basis of complex traits.

## METHODS

### Animal

Adult reissner lamprey (*Lethenteron reissneri*) individuals used in this study were collected from the Songhua River in Heilongjiang Province, China. Specimens ranged from 13 to 16 cm in body length and weighed 10–15 g. The animals were carefully selected and maintained in a temperature-controlled, automated water purification system at 4–6 °C. All experimental procedures were conducted in accordance with Chinese animal welfare legislation.

### Estimation of genome size by flow cytometry

Lampreys were euthanized by immersion in MS-222 (150 μg/ml, Sigma-Aldrich, E10521), and eight tissues (sperm, testis, brain, gill, liver, heart, muscle, intestine, and kidney) were immediately dissociated into single-cell suspensions. Except for sperm, other tissues were cut into small pieces of 0.05 mm and incubated with 0.05 % collagenase I (Sigma-Aldrich, 1148089) at room temperature for 10 minutes. The reaction was stopped with 10 % fetal bovine serum (FBS, Sigma-Aldrich, 12103C) and filtered using 40 μm cell filter for DAPI staining (1 μg/ml, Sigma-Aldrich, D9542).

Flow cytometry was performed on single-cell suspensions from six individuals using a FACSCalibur flow cytometer (BD Biosciences). Nuclear DNA content was estimated by comparison to an internal standard (mouse hepatocyte nuclei) and averaged across four captures of 10,000 events each, temporally interleaved across individuals and tissues.

### DNA isolation and sequencing

All sperms from a healthy and sexually mature male lamprey were collected, and the sperms appeared off white with no blood contamination. Sperms (260 μl, 2.75×10^10^pieces/mL) was divided into three parts and used for long-read sequencing, Illumina sequencing, and Hi-C sequencing.

For long-read sequencing, high-molecular-weight genomic DNA were extracted for 200 μl sperm (2.75×10^10^pieces/mL), followed by purification with Monarch® HMW DNA Extraction Kits (T3060S). Genomic DNA was used to construct both PacBio HiFi and ultra-long ONT sequencing libraries. Quality and quantity of total DNA were evaluated using a NanoDrop One UV‒Vis spectrophotometer (ThermoFisher Scientific, Waltham, MA) and Qubit 3.0 Fluorometer (Invitrogen life Technologies, Carlsbad, CA), respectively, and PFGE (Pulsed Field Gel Electrophoresis, Bio-Rad, CHEF Mapper XA) is used to determine the length of DNA sufficient to obtain longer HiFi and ultra-long ONT reads.

Ultra-long ONT libraries were sequenced on the Nanopore PromethION platform, while PacBio libraries were run on the PacBio Sequel II/IIe system. For ONT ultra-long sequencing, we collected raw sequencing data from 10 flow cells, which generated 235.61 Gb (188.5x) data, among which 139.25 Gb (111.4x) ultra-long reads exceed 100 kb. For genome assembly, only ultra-long ONT reads ≥100 Kb in length were used.

For the PacBio HiFi sequencing (Pacbio Sequel II/IIe platforms), raw subreads were first processed by removing junction sequences and filtered using a minimum length threshold of 50 bp. High-accuracy HiFi reads were then generated using the Circular Consensus Sequencing (CCS, v6.4.0) algorithm with the parameters ‘min-passes=3’ and ‘min-rq=0.99’, ensuring all reads had quality scores above Q20. In total, 122,397,942,993 bases of valid HiFi reads were obtained, with a read N50 length of 16.49 kb.

For Hi-C sequencing, sperm pellets (50 μl, 2.75×10^10^ pieces/mL) were fixed with formaldehyde for DNA cross-linking. After cross-linking was completed, the cells were lysed and then taken for quality testing. Next, the samples were subjected to biotin labeling, blunt-end ligation, and DNA purification to prepare Hi-C samples, and samples proceeded to the standard library construction process. This process included removing biotin from the Hi-C fragments, fragmenting the DNA through sonication, repairing the DNA ends, adding A bases, and ligating sequencing adapters to form adapter-ligated products. PCR condition screening and amplification were then performed to obtain library products. Samples of the library amplification products were taken for Hi-C junction quality control testing. The constructed library was then sequenced using the MGI-2000 sequencer, employing a sequencing strategy of PE150.

For Illumina sequencing, sperms (10 μl, 2.75×10^10^ pieces/mL) were washed with pre-cooled Hank’s buffer, and centrifuged to isolate cells. Cell pellets were lysed with buffers containing 1% SDS, RNase A and Proteinase K, followed by phenol: chloroform: isoamyl alcohol (25:24:1) extraction to remove proteins and other contaminants. DNA was precipitated with isopropanol, washed with 75% ethanol, and resuspended in Buffer EB. DNA quality and yield were evaluated by spectrophotometry and gel electrophoresis. Paired-end and mate-pairs Illumina libraries were generated using Illumina’s paired-end kits according to the manufacturer’s instructions. The libraries were sequenced on an Illumina HiSeq 2500 platform.

### DNA isolation and NGS sequencing of somatic tissues

For the adult lamprey (three samples - gill, heart and intestine), 200 μg of DNA of each sample was sheared into ∼350 bp insert-size fragments using paired-end kits (Illumina, San Diego, CA, USA). All libraries were sequenced using 150-bp paired-end sequencing on an Illumina HiSeq™ 2500. A total of 100 Gb sequenced data was obtained. On average, 30 Gb (30-fold coverage of the genome) of the sequence was obtained. The raw sequenced reads were filtered with SOAPnuke version 1.5.3 with excluding N content (N content >5%), adapter, PCR duplicates, and low-quality (low-quality base >20%, quality value ≤10) reads.

The clean reads were mapped against the new LrT2T genome using BWA-MEM (v0.7.17). Alignment results in bam format were sorted and indexed using samtools (v1.3). *SMC5* genes sequence coverage was identified with IGV (v2.18), to assess whether the gene has been lost in the adult lamprey genome.

### Genome assembly

To estimate the size of the genome, Jellyfish (v2.3.0) was used to count the k-mers by setting the k-mer parameters to 19 and 25 bp, and to obtain the corresponding frequency distributions using the high coverage and quality short reads, which was indexed by applying GenomeScope2 (k-mer =19 bp), with the following parameters: jellyfish count -m 19 -s 100M -t 10 -C Illumina_reads.fasta.

We used hifiasm (v0.20.0-r639) and Verkko (v2.2.1) to assemble the PacBio HiFi, ultra-long ONT (≥100Kb) and Hi-C reads with the following parameters:

hifiasm -o asm -t 64 --h1 HiC_1.fq.gz --h2 HiC_2.fq.gz --ul-cut 100000 --ul ONT.fq.gz HiFi_reads.fasta

verkko -d asm --hifi HiFi_reads.fasta --nano ONT.fq.gz --hic1 HiC_1.fq.gz --hic2 HiC_2.fq.gz.

After comparing the size and continuity of the assemblies, the main assembly (named LrT2T) obtained using Hifiasm was adopted for subsequent analysis (table S4). The initial assembly (LrT2T) yielded 618 contigs, which were polished in three rounds using quality-filtered ONT reads, PacBio HiFi reads, and Illumina reads with NextPolish (v2.2.2) and Pilon (v1.24) with the following parameters:

pilon --genome LrT2T.fasta --fix all --changes --frags LrT2T_illumina.sorted.bam --output pilon --outdir pilon_result --vcf

nextPolish test_data/run.cfg (task = 5, rewrite = no, lgs_options = -min_read_len 5k -max_depth 100, lgs_minimap2_options = -x map-ont / map-pb).

Contigs from those assemblies were used to fill gaps in the draft assembly. Subsequently, the genome was gap filled using TGS GapCloser (v1.2.1) with HiFi and ONT reads, following recommended parameters: tgsgapcloser --scaff HiFi and ONT Reads.fasta --reads T2T.fasta --min_idy 0.1 --min_match 10 --thread 64 --max_candidate 1 --output test_ne --ne. We used the assembled hifiasm version as the primary contig framework and generated assembled Verkko version to verify hifiasm results, patch gaps, and telomere regions where high-quality consensus sequences had better read-to-assembly support by minimap2 (v2.24) and IGV (v2.18). The Graph Manual Curation (Bandagev0.9.0 visualization of the assembled graph generated by Verkko) from the Verkko results was used to verify the gap-filling performed by TGS-Gapcloser. Meanwhile, we used the raw reads to examine the regions with low coverage of the HiFi and ONT reads to ensure that these areas were not caused by errors in the gap filling process performed by TGS GapCloser. We also used Verkko to re-assemble the genome and manually correct any errors that may have been caused by TGS GapCloser, in order to ensure the accuracy of the genome sequence. After gap filling, manually resolved gap sequences and their ± 5 Kb flanking regions were extracted and iteratively polished.

For Hi-C read assignment, we required each read pair to map without mismatches and to be aligned to the 618 assembled contigs. These assigned pairs were used for scaffolding with the 3D-DNA pipeline (v180922) (75). Hi-C reads were aligned to the haploid genome using the Juicer pipeline (v1.7.6), and the resulting alignment information was used by 3D-DNA to generate the Hi-C contact map with the following parameters:

juicer.sh -z LrT2T_pilon.fasta -p genome.chrom.sizes -y genome_DpnII.txt -s DpnII -d ./work/ -D ./juicer -t 32 -S final

run-asm-pipeline.sh -r 0 LrT2T_pilon.fasta merged_nodups.txt.

The contact map was then visualized and manually curated in Juicebox (v2.15), allowing correction of misassemblies (e.g., inversions) and re-joining of unlinked contigs. A refined chromosome-level assembly was then generated by rerunning the 3D-DNA pipeline.

### Assembly quality assessment

To evaluate assembly quality (Contig N50, quality value - QV, and coverage, etc), we first mapped the assembled genome to the published sea lamprey reference using QUAST (v5.2.0) with the following parameters: quast.py contigs.fasta -r reference.fasta -g genome.gff -o quast_out -t 4.

In addition, Merqury (v1.3) was employed to calculate QV using Pacbio HiFi reads with the following parameters: merqury.sh LrT2T_k19.meryl LrT2T.fasta out_prefix. The LrT2T_k19.meryl databases (k-mer =19 bp) were generated from PacBio HiFi reads separately using Meryl (v1.4.1), and k-mers that appeared only once were filtered out. Finally, QVs were calculated using Merqury with default settings, following the formulas: QV = −10log_10_(error k-mer counts/total k-mer counts).

Genome completeness was further assessed using BUSCO (v5.7.0, parameters: run_BUSCO.py -i LrT2T.fasta -c 5 -o busco_LrT2T -m genome -l metazoa_odb10 --offline) with the metazoan dataset (metazoa_odb10), which includes 954 universal single-copy orthologs. In addition to the reference genome described in the main text, the genomes of other lampreys (*Petromyzon marinus*-GCF_010993605.1, *Entosphenus tridentatus*-GCA_014621495.2, *Lampetra auremensis*-GCA_029874635.1, *Lethenteron camtschaticum*-GCA_018977245.1, *Lampetra planeri*-GCA_965212315.1 and *Lampetra fluviatilis*-GCA_964198595.1) were used for quality assessment and comparison.

### Identification of centromeres and telomeres

Telomeric and centromeric repeat sequences in the assembled genome were identified using CentIER (v2.0), TRASH (v1.0) and TIDK (v0.2.1), based on the distribution patterns of repetitive elements along chromosomes.

For centromere identification, we applied CentIER (v2.0) (https://github.com/simon19891216/CentIER) and TRASH (https://github.com/vlothec/TRASH) with the following parameters:

centIER.py --matrix1 example1_100000.matrix --matrix2 example1_20000.matrix --bed1 example1_100000_abs.bed --bed2 example1_20000_abs.bed --gff T2T.gff3 T2T.fasta

TRASH_run.sh assembly.fa --o output.path --t template_file.csv --horclass classname.

Rather than relying solely on tandem repeat detection, we employed an integrative approach that considers multiple sequence features, including tandem repeat density, retrotransposon content, and k-mer frequency distributions. The putative centromeric regions were further defined using genome structural annotation data as input.

The output results of CentIER and TRASH were scored for each sequence at a 1:1 ratio (with a total score of 10). Sequences with a score of 5 or higher are considered to be putative centromere sequences. We used the inferred chromosome centromere sequence and the centromere region of each chromosome for blast, and determined the distribution of repetitive motifs in the potential centromere regions of 90 chromosomes (data S2). The types of monomer repeats were determined based on the sequence characteristics of the putative centromere regions (1-10 Mb) of each chromosome, and they were searched for homology in all chromosomes using a threshold of 70%. We have identified a 48 bp repeat unit (named Cen-3), which was used to search for similar repeats that contain homologous sequences (including continuous GT repeats) in the centromeric regions of other chromosomes. Cen-3 turned out to be the most common sequence motifs in the inferred centromere regions of all chromosomes (78.89%).

Notably, we observed that a large number of atypical repetitive sequences were enriched at both chromosomal ends (∼1% of the chromosome end regions), indicating potential telomere-associated regions in the lamprey genome. To refine the identification of telomeric repeat units, we employed TIDK with the following parameters:

tidk explore -f genome.fa --minimum 5 --maximum 12 -o tidk_explore -t 2 --log --dir telomere_find --extension TSV.

This analysis yielded a set of repeat units designated as telomere-associated sequences, with detailed sequences provided in data S3.

### Fluorescent in situ hybridization (FISH) for centromeric and telomeric repeats

Lampreys were intraperitoneally injected with colchicine (2 μg/g body weight) and maintained at 4 °C for 16 hours. Fresh testes were then collected, dissociated into a single-cell suspension, and fixed with Carnoy’s solution (ethanol: glacial acetic acid, 3:1, v/v). Amplification of the centromeric (Cen-3, 48bp) and telomeric (Tel-104) repeat sequences was performed using specific primers listed in Datas S2 and S3.

Cen-3 sequence:

5’-GTGTGTGTGGGGGGGGGGGTCACAAGACATGTGGACCACCGCACGCAC-3’

Tel-104 sequence:

5’-ACTCCACTCCACTCCACTCC-3’

The purified PCR products of Cen-3 and Tel-104 were labeled with Cy5-dUTP and FITC-dUTP, respectively, using a Nick Translation Mix (Roche, Mannheim, Germany). Nick translation was carried out at 15 °C for 1.5 hours, and probe quality was verified by agarose gel electrophoresis. Chromosome spreads were prepared following previously described protocols (76), with one optimization: the hypotonic treatment step was performed using 0.005% sodium citrate solution.

Slides containing metaphase cells were selected for the DNA FISH assay. Chromosome preparations were denatured in 70% formamide for 1 minute at 70 °C. Each 20 μL hybridization mixture contained 50% formamide, 10 mg/mL dextran sulfate, 2× SSC, 100 μg/mL DAPI (62248, Thermo Fisher Scientific), and 100 ng of each fluorescently labeled probe. Hybridization was carried out at 37 °C for 5 hours. Following hybridization, slides were washed three times in 1× SSC for 5 minutes each and rinsed in purified water at room temperature. Slides were then air-dried in the dark. Fluorescence images were captured using a laser scanning confocal microscope (Carl Zeiss LSM 710) and processed using Adobe Photoshop (v19.0, Adobe Systems, San Jose, CA, USA).

### RNA-seq data generation and genome annotation

To generate RNA sequencing data for gene annotation, total RNA was isolated from the embryos (4dpf, 6dpf, 8dpf and 12dpf), larva (16dpf, 20dpf, 25dpf and 30dpf), brain, liver, kidney, intestines, muscle, supraneural, gill and heart of a male individual using the EasyPure RNA Kit (Transgen). Sequencing libraries were generated using the NEBNext® UltraTM RNA Library Prep Kit for Illumina® (NEB, Ipswich, MA, USA), following the manufacturer’s recommendations. A total of 600Gb RNA seq data and 1,100Gb Iso-seq (long-read sequencing) data were collected for assisting genome annotation (table S2).

The first step in the workflow involves purifying the poly-A containing mRNA molecules using poly-T oligo-attached magnetic beads. Following purification, the mRNA is fragmented into small pieces using divalent cations under elevated temperature. The fragmented RNA is then reverse transcribed into first-strand cDNA using reverse transcriptase and random primers. Second-strand cDNA synthesis is subsequently performed using DNA Polymerase I and RNase H. After end repair and the addition of a single ‘A’ base, sequencing adapters are ligated to the cDNA fragments. The resulting products are purified and enriched via PCR amplification. PCR yield is quantified using Qubit, and the samples are pooled and converted into single-stranded DNA circles (ssDNA circles) to generate the final sequencing library.

DNA nanoballs (DNBs) were generated with the ssDNA circle by rolling circle replication (RCR) to enlarge the fluorescent signals at the sequencing process. The DNBs were loaded into the patterned nanoarrays and pair-end read of 125 bp or 150 bp were read through on the BGISEQ-500 platform for the following data analysis study. For this step, the BGISEQ-500 platform combines the DNA nanoball-based nanoarrays and stepwise sequencing using Combinational Probe-Anchor Synthesis Sequencing Method.

HiSat2 (v2.1.0) was used to align the RNA-seq reads to the genome of Lethenteron reissneri. StringTie (v2.0) was used for assembling the transcripts and estimating abundances. We also used the gene annotation of this assembly. The NOISeq R package (v2.31.0) was used for differential expression tests.

For Iso-seq sequencing, the library was constructed as per Clontech SMARTer-PCR cDNA Synthesis Sample Preparation Guide”. 1–2 Kb libraries were selected on 0.8% Agarose Gel, purified, end repaired and blunt end SMARTbell adapters were ligated. The libraries were quantified using Qubit (Invitrogen) and validated for quality by running a LabChipGX (Caliper Life Sciences) subsequently sequencing was done in 8-well-SMART Cell (v3) in PacBioRSII. The generated raw sequences were processed for quality control checks including filtering of high-quality reads based on the score value, removal of primer/adaptor sequence reads and trimming of read-length using SMRTLink (v8.0).

Our process has drawn upon the genetic prediction pipeline of the sea lamprey, including 1) Genome annotations were produced using the Augustus GeneMarkS-T and SNAP and other genome annotation pipeline. 2) Additional input included previously published mRNA-seq reads derived from lamprey embryos. 3) dentification pipeline, and putative gene functions were assigned using BLASTP identified homology to the Uniprot/Swiss-Prot protein database (data S1).

The gene prediction pipeline of LrT2T was expanded by adding a large number of mRNA-seq reads and Iso-seq reads of other tissues, and putative gene functions were assigned using more extensive database (e.g., Gene Ontology, KEGG, Swissprot, NR, TrEMBL, EGGNOG, InterPro, and IPR domains) (data S1). Notably, we generated 1,100Gb full-length transcript Iso-seq and 800Gb RNA-seq data across eight tissues and four developmental stages. The great majority (95.78%) of the annotated genes were based on homology prediction and RNA sequencing data (table S2). These optimizations have made the genome annotation of the LrT2T gene more reliable, and we used the RNA-seq reads of the reissner lamprey and the sea lamprey for mutual annotation to further improve it.

For genome annotation, the predicted repeat families and tandem repeats were combined with the existing fish and agnatha repeat library from Dfam (v3.1) and RepBase (v20170127) as the input library for repeat annotation and masking with the program RepeatMasker (v4.0.7). We used the braker program to translate the existing RefSeq gene model annotations into the new assemblies. RNA-seq-based gene prediction was performed by mapping clean RNA-seq reads to the genome using Hisat (v2.1.0) and assembled by StringTie (v2.0). Multiple methods including PASA (v2.0.2), TransDecoder (v.2.0) and GeneMarkS-T (v.5.1) were applied to predict coding regions. AUGUSTUS (v.2.4), GlimmerHMM (v.3.0.4), GeneID (v.1.4) and SNAP were used for de novo gene prediction with default parameters. UniGenes were then inputted to PASA (v2.0.2) to predict genes. Training models used in AUGUSTUS, Glimmer HMM and SNAP were obtained from the prediction results of PASA and GeMoMa (v1.3.1). Gene models from these different approaches were combined by EVM (v1.1.1).

The predicted genes were annotated by blasting their sequences against a number of nucleotide and protein sequence databases, including COG Release, KEGG Release, NCBI NR Release and SWISS-PROT Release. Moreover, these predicted genes were annotated against the Pfam database of the HMMER (v3.1b2) software (http://www.hmmer.org) and the InterPro database of InterProScan (v5.34) (https://github.com/ebi-pf-team/interproscan). Gene Ontology for each gene was assigned by Blast2GO (v2.5) based on NCBI databases.

### Gene expression time-series RNA-seq data with Mfuzz

Initially, using the one-way analysis of variance (ANOVA), genes significantly differentially expressed across eight sample groups (4dpf, 8dpf, 12dpf, 14dpf, 16dpf, 20dpf, 25dpf and 30dpf) were identified. Following this, the Mfuzz (v2.29) package was utilized for clustering analysis on the expression patterns of these identified genes according to their sequential order through the stages. The genes exhibiting synchronous changes in expression patterns throughout all stages, which reflect embryos and larva development process for lamprey, were then identified. These factors, inferred to be closely associated with heart development and chamber formation, were incorporated into subsequent analyses aimed at constructing the correlation between *CDH2_H* and cardiac development and chamber formation.

### Comparative genomic analyses and identification lineage-specific expansions of gene families

Gene structures in the reissner lamprey genome were annotated using GeMoMa (v1.4.2) with default parameters, based on available gene models. Predictions were refined by applying a GeMoMa annotation filter with default parameters, except for the evidence percentage threshold, which was set to e = 0.1. Predicted transcripts were then manually curated to retain a single high-confidence transcript per locus.

Homology analysis was conducted using OrthoFinder (v2.5.2, orthofinder -f ./Orthofinder/pro -S diamond - n pro -t 64 -a 64 -p $TMPDIR) across the LrT2T genome and other representative vertebrate reference genomes. Protein sequences corresponding to the longest and most reliable transcript per gene in the LrT2T genome were used. A gene was considered homologous to those in the NAM founder lines if its protein sequence belonged to the same orthogroup as at least one NAM gene according to OrthoFinder. Additionally, protein sequences from the LrT2T genome were aligned to those from the sea lamprey (NCBI v3) using BLASTP (v2.9.0) with an E-value cutoff of <1 × 10^−3^ to further confirm homology.

For gene family expansion analysis, using OrthoFinder with an all-vs-all BLASTP search (E-value < 1 × 10^−5^) and Markov clustering (inflation parameter = 1.5), we first identified orthologous genes and their protein sequences from the high-quality genomes of 75 species, and they were compiled and clustered into 30,098 orthogroups. These 75 species cover the major vertebrate lineages, including *Ailuropoda melanoleuca*(GCA_002007445.3), *Analcellicampa danfengensis* (GCF_028878055.3), *Anas platyrhynchos* (GCA_047663525.1), Anolis carolinensis (GCA_035594765.1), *Balaenoptera musculus* (GCA_009873245.3), *Bos taurus* (GCA_002263795.4), *Branchiostoma belcheri* (GCA_001625305.1), *Branchiostoma floridae* (GCA_000003815.2), *Branchiostoma lanceolatum* (GCA_035083965.1), *Callorhinchus milii* (GCA_018977255.1), *Canis lupus familiaris* (GCA_011100685.1), *Carcharodon carcharias* (GCA_017639515.1), *Chelonoidis abingdonii* (GCA_003597395.2), *Chelydra serpentina* (GCA_037349315.2), *Chrysemys picta bellii* (GCA_011386835.2), *Chrysolophus pictus* (GCA_037305975.1), *Ciona intestinalis* (GCA_000224145.2), *Ciona savignyi* (GCA_000149265.1), *Coturnix japonica* (GCA_001577835.2), *Crocodylus porosus* (GCA_001723895.1), *Delphinapterus leucas* (GCA_002288925.3), *Equus caballus* (GCA_041296265.1), *Eptatretus atami* (GCA_035128595.1), *Eptatretus okinoseanus* (GCA_041222605.1), *Felis catus* (GCA_018350175.1), *Gambusia affinis* (GCA_019740435.1), *Hemiscyllium ocellatum* (GCA_020745735.1), *Hemitrygon akajei* (GCA_048418815.1), *Lampetra richardsoni* (GCA_050491395.1), *Lepisosteus oculatus* (GCA_040954835.1), *Lethenteron camtschaticum* (GCA_018977245.1), *Leucoraja erinaceu* (GCF_028641065.1), *Loxodonta africana* (GCA_030014295.1), *Macaca mulatta* (GCA_003339765.3), *Mastacembelus armatus* (GCA_900324485.3), *Meleagris gallopavo* (GCA_000146605.4), *Monopterus albus* (GCA_001952655.1), *Mordacia mordax* (GCA_036362905.1), *Naja naja* (GCA_009733165.1), *Neogobius melanostomus* (GCA_007210695.1), *Oncorhynchus mykiss* (GCA_013265735.3), *Ornithorhynchus anatinus* (GCA_004115215.4), *Oryctolagus cuniculus* (GCA_964237555.1), *Ovis aries* (GCA_016772045.2), *Pavo cristatus* (GCA_045791835.1), *Pelodiscus sinensis* (GCA_049634645.1), *Phasianus colchicus* (GCA_004143745.1), *Physeter catodon* (GCA_002837175.5), *Pogona vitticeps* (GCA_047335585.1), *Pongo abelii* (GCA_028885655.3), *Pygocentrus nattereri* (GCA_015220715.1), *Scophthalmus maximus* (GCA_022379125.1), *Sus scrofa* (GCA_000003025.6), *Taeniopygia guttata* (GCA_048771995.1), *Takifugu rubripes* (GCA_901000725.3), *Tetraodon nigroviridis* (GCA_051020865.1), *Tursiops truncatus* (GCA_011762595.2), *Varanus komodoensis* (GCA_004798865.1), *Zalophus californianus* (GCA_009762305.2), and reissner lamprey (LrT2T).

To identify lineage-specific gene family expansions, orthogroups were analyzed using OrthoFinder’s gene duplication inference in conjunction with a species phylogeny visualized via iTOL v6 (https://itol.embl.de/). To examine expansions specific to the vertebrate common ancestor, 1,562 single-copy orthologues from the ten species were extracted and subjected to multiple sequence alignment using MUSCLE v3.8.31. Gene Ontology (GO) enrichment analysis was performed in R using the Database for Annotation, Visualization and Integrated Discovery (DAVID). Gene-Concept network was generated using the 69 genes with vertebrate-specific expansion and enriched in cardiac-related GO terms by cnetplot and Cytoscape (v3.9.1). The *CDH2* gene tree was constructed using MEGA (v10.1) with 1,000 bootstrap replications. The LG substitution model, and the Nearest Neighbor-Interchange heuristic model were used.

### RNA-seq and single-cell RNA-seq data analysis

After trimming based on quality scores using Btrim (v0.2.0), clean reads were aligned to the LrT2T genome with TopHat (v2.1.1). Gene abundance in different tissues was calculated using Cufflinks (v2.1.1).

Single-cell RNA-seq data was obtained from the published study (42), which 604,460 cells/nuclei and 70 cell types from 14 lamprey tissue samples (PRJNA1194219).

The *CDH2* expression data of representative vertebrates used in Figure 4H, except for the lamprey, were obtained from the publica databases (human - https://data.humancellatlas.org/, mouse - https://bio.liclab.net/SCInter/index.php, zebrafish - https://singlecell.broadinstitute.org/single_cell/study/SCP162/ and lancelet - https://lifeomics.shinyapps.io/shinyappmulti/).

### Luciferase reporter gene assay

The CRE elements (cis-regulatory elements) of *CDH2* in higher vertebrates was obtained using UCSC Genome Browser (http://genome.ucsc.edu/index.html), and confirm that the CRE elements is conserved in zebrafish, mice, and humans. The enhancer sequence was used for genome blast in lampreys and lancelet. The conserved enhancer sequence of the vertebrate *CDH2* gene has been identified as appearing in the lamprey *CDH2_H* gene. The enhancer sequence (338bp) is:

5’TTATGAGAGAATGTGATCAATATGTTTGTTGATCTTGAAAATAGCCTAGGAAAGTGAAAGGATAGATTAATTAAAGAATTAAGCCAAACATGAATTTTAATATTCAGAGACATTTATAAGCAAAACACCCTAAGGTATATTTATAGAGCCACTTTAATTAGGCAGTATTCAAAACAGCTTTACACATGAGTCAAGTTTTCAACGTACTTTAAATATGCATCTCTTCTAAATGAGAGAGCTGGGGTATTGTAAACCAAGCATGGAACCTGGAATCAGAACATGAGGCCTTGAGTCCTGGCTCTGTGGGTTACAAACTCTGAAAGTAAATCATGCAACTT3’.

P1 (postnatal day 1) mice were anesthetized on ice, and their hearts were collected with scissors and forceps. The 20 neonatal mouse hearts were removed in PBS. The hearts were added to a C-tube (130-093-237, Miltenyi, Germany) with the Neonatal Heart Dissociation Kit (2.5 mL of enzyme mixture). The hearts in the C-tube were then placed in a gentleMACS Octo Dissociator (130-095-937, Miltenyi, Germany). The digestion time was approximately 2 min. At the end of the procedure, the digestion was immediately stopped by adding 5 ml FBS. The cell suspension was filtered with a 100-μm strainer and centrifuged at 300 ×g for 5 min. The supernatant was aspirated carefully. Subsequently, the isolated cardiac cells were plated on culture flasks with DMEM containing 10% FBS at 37 °C and 5% CO_2_ for 90 min to remove the attached non-cardiomyocytes, and the cell suspension containing mainly cardiomyocytes was then harvested.

Enhancer expression plasmid or its control plasmid (i.e., PGL3-promoter vector) was co-transfected into primary cultured mouse cardiomyocytes (PCMCM) with a single report plasmid by Lipofectamine TM 3000 (Thermo Scientific, L3000001). If the 338bp sequence has enhancer activity, the transfected cell line exhibits firefly luciferase activity. Luciferase was measured 48 h after transfection by ultrasonic lysis of cells. The firefly luciferase activity was then normalized to β-galactosidase activity. Experiments were repeated at least three times.

### Lamprey spawning and husbandry

Eggs and sperm were manually stripped from ripe adult lampreys during the spring spawning season. For *CDH2_H* and enhancer gRNA injections, ten females were used across three independent fertilization events. Sperm from five males were pooled for each fertilization. Embryos and larvae were maintained at 18 °C in individual dishes containing 10 L of deionized water supplemented with 400–600 ppm artificial sea salt. Water was changed daily, and dead embryos were removed to prevent contamination. Developmental staging was performed according to Nikitina et al. (77). Injected embryos (including negative controls) exhibited a slight developmental delay; therefore, staging was based on morphological landmarks of unaffected embryos rather than chronological age, typically corresponding to a 0.5–1 day delay relative to wild-type siblings.

### CRISPR/Cas9-mediated CDH2_H and enhancer knockout

The CRISPR/Cas9 gene-editing system was employed to generate *CDH2_H* and enhancer knockouts in lamprey. Three target sites were selected within the first exon of *CDH2_H* or within the 338 bp enhancer region. Target sites were chosen based on the following criteria: (1) GC content between 50 % and 80 %; (2) proximity to the presumed start codon; and (3) absence of off-target matches in the available sea lamprey or reissner lamprey genomes, defined as having > 80 % similarity by BLAST or fewer than three mismatches within the ten bases proximal to the PAM sequence. The sequences of the sgRNAs targeting *CDH2_H* or the enhancer are listed in table S8. The control sgRNA was designed according to the published study (78) (table S8).

The sgRNAs (chemically synthesized; GenScript, HK.1548; concentration: 20 ng/μL each) were co-injected with SpCas9 protein (GenePharma, Suzhou, China; final concentration: 300 ng/μL) into one-cell-stage fertilized eggs. Microinjections were performed in three independent experiments, each consisting of three groups: *CDH2_H* knockout (*CDH2_H*-ko), enhancer knockout (enhancer-ko), and control (negative control sgRNA were co-injected with SpCas9 protein) (table S8) (78). For each group, 4,000 fertilized eggs were injected, and the experiments were done in three replicates, resulting in a total of 12,000 embryos for each group.

To assess knockout efficiency, embryos surviving to 6 dpf and 12 dpf were pooled (n=50 per group) for total DNA extraction and subjected to 50× Illumina whole-genome sequencing. BAM files generated from genome alignment were analyzed using IGV to evaluate editing efficiency. Additionally, embryos used for assessing cardiac development at 12 dpf were individually genotyped via Illumina sequencing using both forward and reverse primers (data S7, data S8).

### Real-time quantitative PCR (RT-qPCR)

Quantitative PCR (qPCR) experiments were performed using the SYBR® PrimeScript™ RT-PCR Kit (TaKaRa, China) according to the manufacturer’s instructions, including genomic DNA removal and reverse transcription. Each reaction contained 1× SYBR Premix Ex Taq, 10 μM of each primer, and 2 μL cDNA (50 ng/μL) in a final volume of 25 μL. Amplification was conducted on a PCR Thermal Cycler Dice Real-Time System (TaKaRa, China) under the following conditions: initial denaturation at 95 °C for 30 s to activate the DNA polymerase, followed by 40 cycles of 95 °C for 5 s, 60 °C for 30 s, and 72 °C for 30 s.

Primers were designed to target exon 1 of each gene, and the specific primer sequences are listed in table S8. *GAPDH* (glyceraldehyde-3-phosphate dehydrogenase) was used as an internal control. Each sample was analyzed in triplicate using the Thermal Cycler Dice Real-Time System analysis software (TaKaRa, China). The specificity of amplification was verified by melting curve analysis.

### Image acquisition and statistical analysis

Lamprey larvae were analyzed using a fluorescence microscope (SMZ1500, Nikon, Japan) and photographed with a digital camera. Quantitative image analyses were performed using NIS-Elements D3.1 (Nikon, Japan) and ImageJ software (National Institutes of Health, Bethesda, MD, USA). Statistical analyses and graphical representations were conducted using GraphPad Prism 8.0 (GraphPad Software, San Diego, CA, USA). P-values were calculated using ANOVA, as implemented in GraphPad Prism.

### Morphological and functional analyses of the lamprey heart

Heart tissues from adult (5-year-old) and larval (12 dpf) lampreys were processed for morphological and ultrastructural examination using hematoxylin and eosin (HE) staining and transmission electron microscopy. Frozen sections were thawed, dewaxed in xylene, and rehydrated through a graded ethanol series. Hematoxylin staining was performed for 10 minutes, followed by dehydration through graded ethanol and xylene. Finally, the sections were mounted with neutral balsam for microscopic analysis. Enhanced hematoxylin staining was used to display myocardial fibers, and after 48 hours of HE staining, dehydration and sealing were performed. The myocardial fibers were stained blue purple.

The heart of 12 dpf lampreys was clearly visible under a stereomicroscope, and heart function was evaluated by counting beats per minute. Still images and videos were captured using the microscope. Cardiac contractility was assessed as previously described (79, 80). Fractional shortening (FS) was determined by measuring the ventricular diameter at the end of diastole and systole using zebraFS software (http://www.benegfx.de). FS was calculated using the formula: FS = (∅diastole − ∅systole) / ∅diastole.

For biochemical analysis, heart tissues from 12 dpf lampreys were dissected under a stereomicroscope. The levels of LDH (A020-2-2), AKP (A059-2-2), AST (C010-1-1), and ALT (C009-2-1) were measured using commercial assay kits (Nanjing Jiancheng Bioengineering Research Institute, China). Approximately 100 mg of lamprey tissue was mechanically homogenized in 1 mL of the corresponding lysis buffer. All enzyme activity measurements were normalized to the total protein content of each sample.

## RESOURCE AVAILABILITY

### Lead contact

Requests for further information and resources should be directed to and will be fulfilled by the lead contact, Bing Su.

### Materials availability

This study did not generate new unique reagents.

### Data and code availability

The genome assembly and annotation data for LrT2T are available under BioProject accession number PRJCA042445 at the Genome Warehouse (https://ngdc.cncb.ac.cn/gwh/). The raw sequencing data, including PacBio HiFi, Oxford Nanopore (ONT), Hi-C, Illumina whole-genome sequencing, and RNA sequencing, have also been deposited under the same BioProject and BioSample accession number SAMC5594263. Processed single-cell RNA-seq data and accompanying metadata are available from the Zenodo repository at https://doi.org/10.5281/zenodo.14338297. All other data supporting the findings of this study are provided in the main text or supplementary materials.

## ACKNOWLEDGMENTS

We thank Yan Guo for her technical assistance in this study.

This study was supported by grants from National Natural Science Foundation of China (32230014 to B.S and Y.P, 32470558 to Q.L), Key Basic Research Project of Yunnan Province of China (202401BC070014 to B.S), Yunnan Scientist Workshop Project (to B.S.), National Science and Technology Innovation 2030 Major Program (STI2030-2021ZD0200100 to B.S), and State Key Laboratory of Genetic Resources and Evolution (GREKF22-14 to Y.P).

## AUTHOR CONTRIBUTIONS

Conceptualization: QL, YP, BS

Methodology: ZD, KL, YH

Investigation: ZD, HW, JL, LL, JL, CZ

Visualization: ZD, HW

Funding acquisition: QL, YP, BS

Project administration: YP, ZD

Supervision: BS, YP, QL, YH

Writing – original draft: ZD, HW, BS

Writing – review & editing: BS, HW, ZD, YP, QL

## DECLARATION OF INTERESTS

Authors declare that they have no competing interests.

## DECLARATION OF GENERATIVE AI AND AI-ASSISTED TECHNOLOGIES

The authors conducted the research and writing process without employing any generative AI tools or AI-assisted technologies.

## SUPPLEMENTAL INFORMATION

Figures S1–S16, and Tables S1–S8 Data S1-S8.

**Figure S1.**
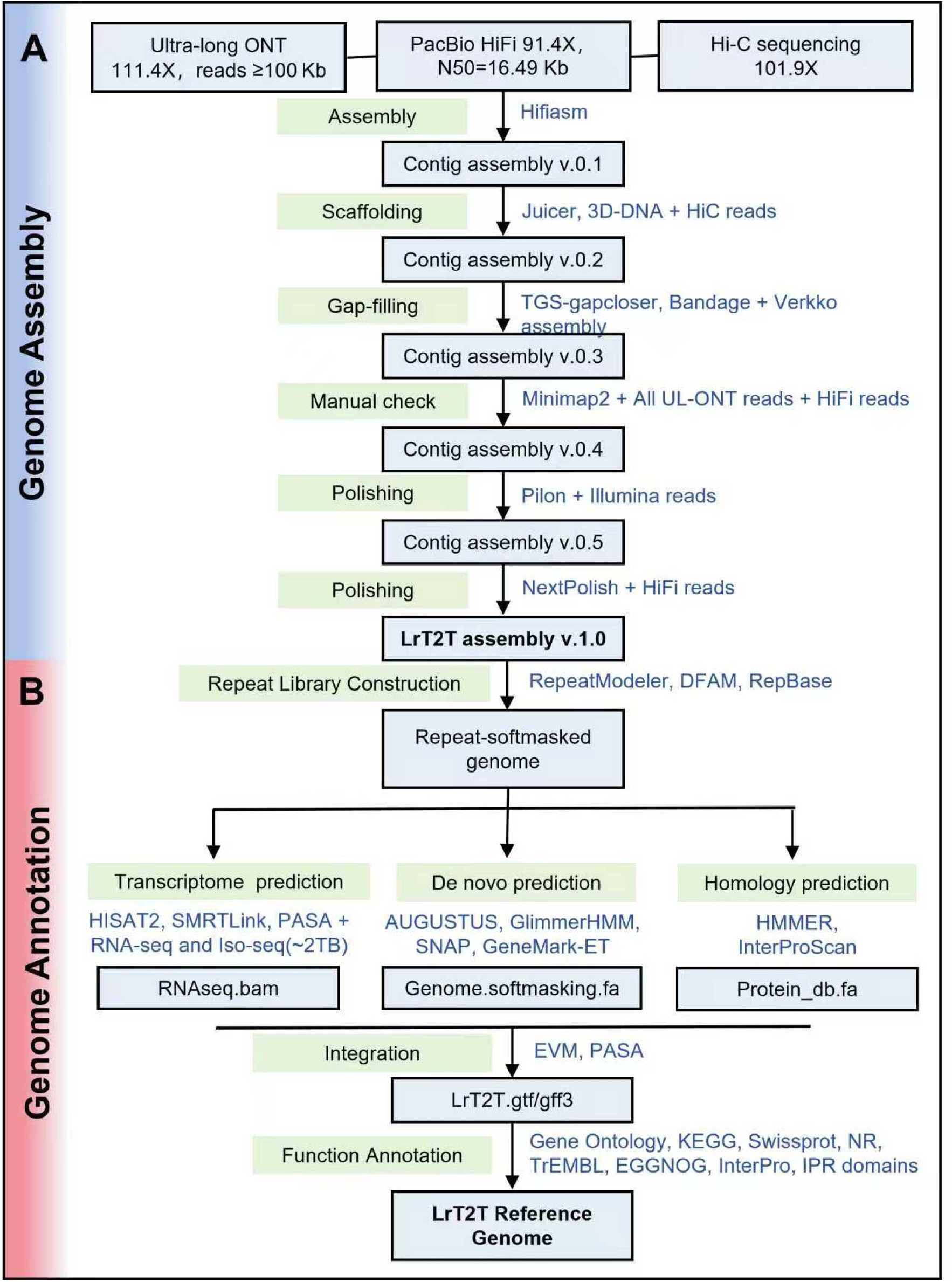
Workflow for telomere-to-telomere assembly and annotation of the LrT2T genome. A schematic representation of the LrT2T genome assembly strategy, encompassing primary assembly, scaffolding, gap filling, polishing and downstream structural/functional annotation. All software and algorithms implemented at each step are highlighted in blue. Detailed procedures are provided in Methods.

**Figure S2.**
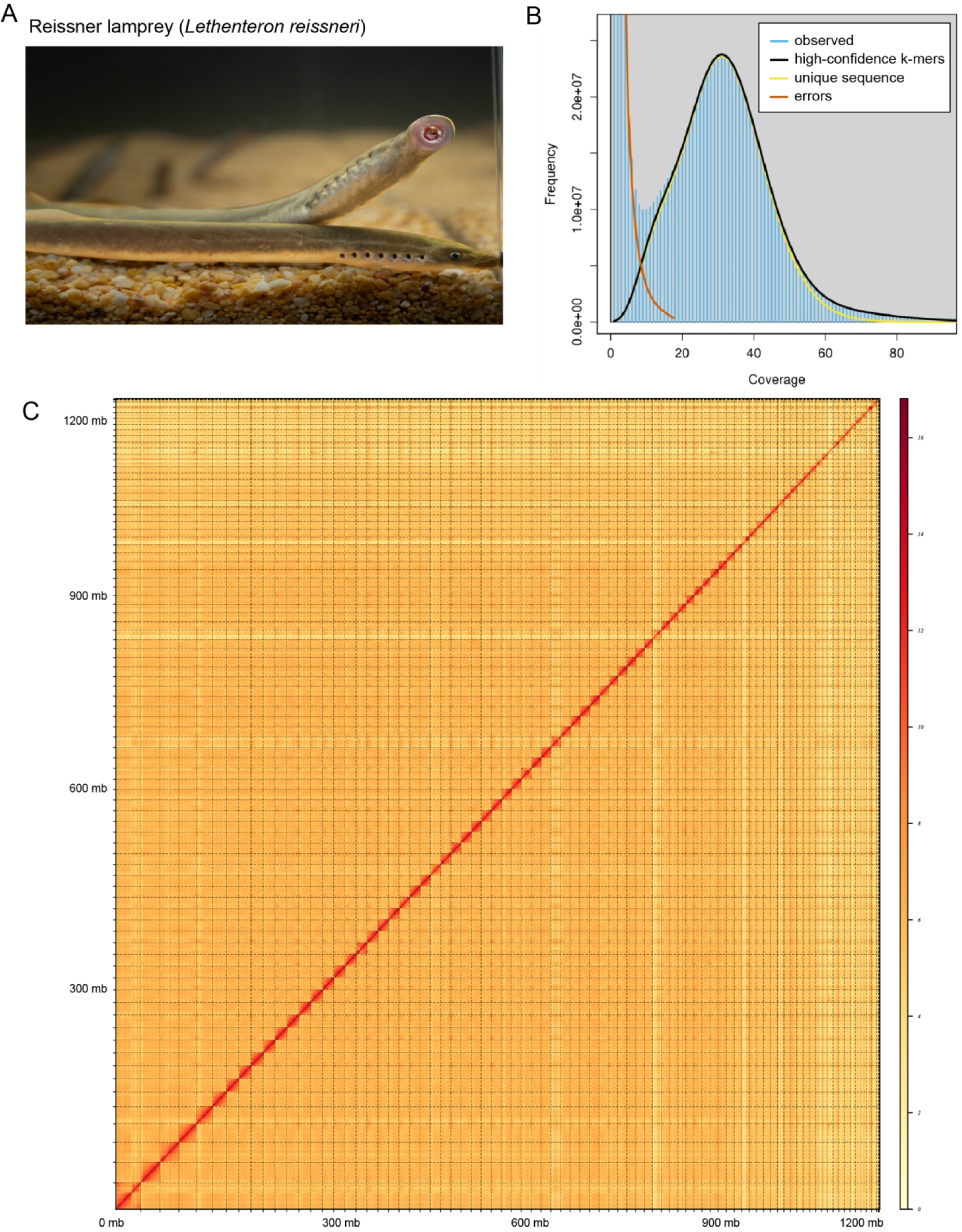
Photograph of the reissner lamprey and k-mer analysis of Illumina sequencing reads. (**A**) Photograph of the reissner lamprey (*Lethenteron reissneri*). (**B**) Distribution of k-mer coverage based on the Illumina reads using a k-mer size of 19 bp. The estimated parameters are: k-mer coverage = 17.3. Unique sequence ratio = 65.6%, repetitive sequence ratio = 2.02%, heterozygosity = 0.477%, and error rate = 0.488%. (**C**) Linked density histogram of the assembled scaffolds of the reissner lamprey genome. Blue-lined squares indicate assembled scaffolds corresponding to the 90 chromosomes.

**Figure S3.**
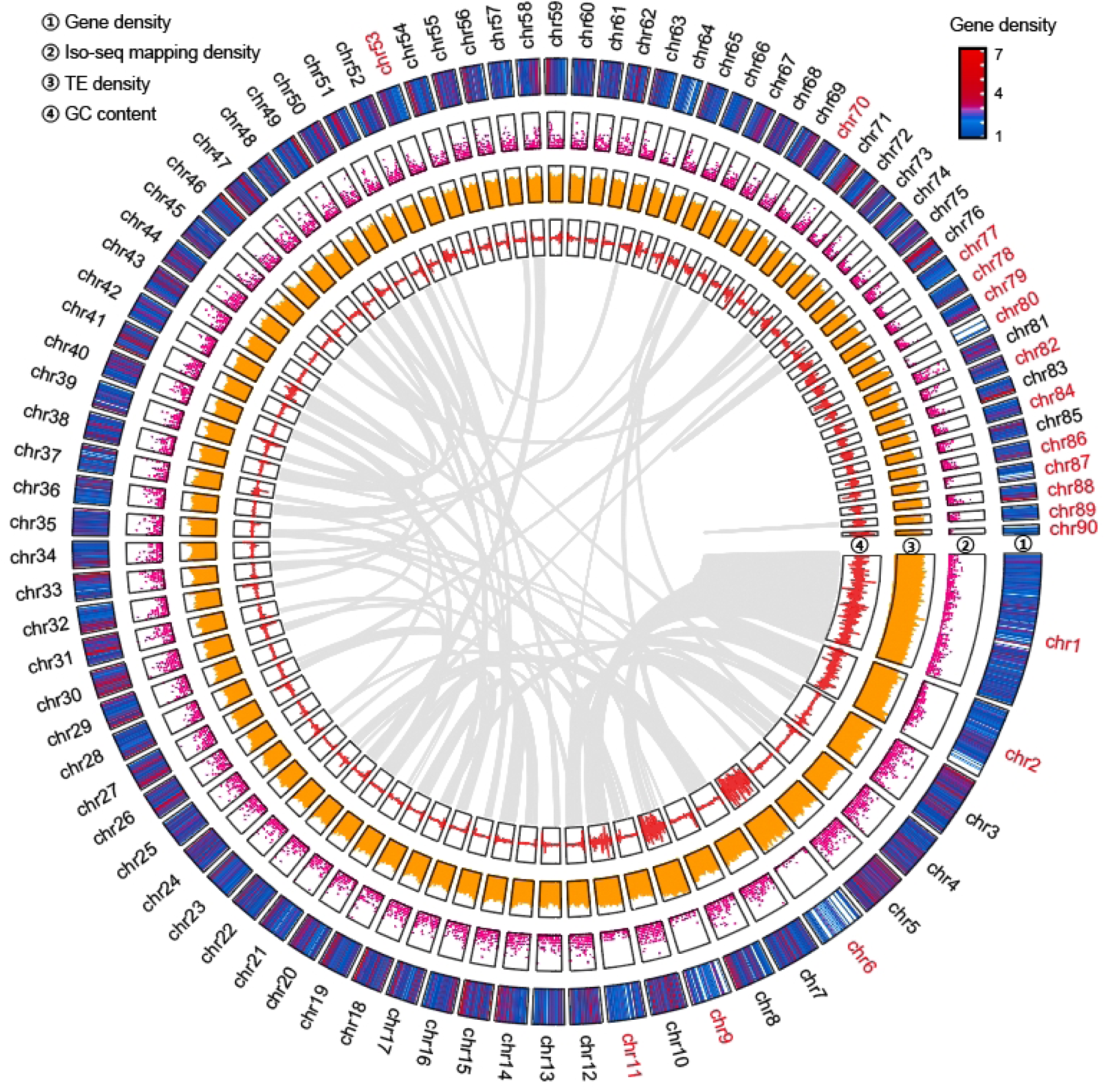
A circos plot of 90 chromosomes of reissner lamprey genome assembly. Overview of the assembled genome showing gene density, Iso-seq mapping coverage, transposable element (TE) density, and GC content across 90 chromosomes. Chromosomes newly assembled to completion in this study (18 chromosomes) are labeled in red. Inter- and intra-chromosomal collinearity is illustrated by gray lines.

**Figure S4.**
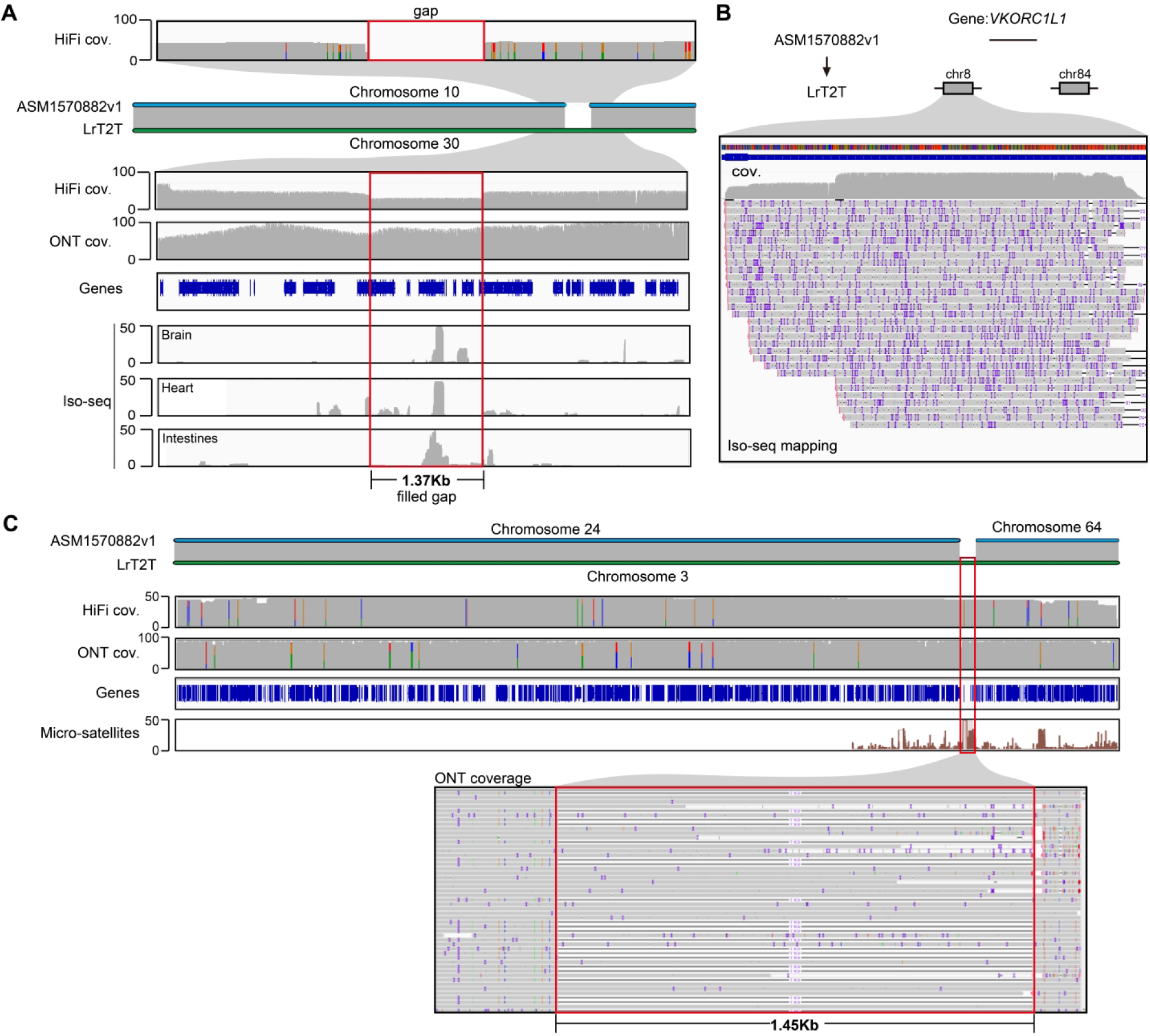
Two examples of assembly gaps resolved in the LrT2T genome. (**A**) A genomic gap (1.37kb in length) in the previous assembly (ASM1570882v1) is resolved in the LrT2T genome. The gap, located on chromosome 10 (chr10) in the previous assembly, showed relatively low coverage by PacBio HiFi reads. In contrast, the LrT2T genome exhibits much higher long-read coverage, particularly from ONT ultra-long reads. Several genes are located within this region, and Iso-seq data confirm their expression across multiple tissues. (**B**) The *VKORC1L1* gene, absent in the previous assembly is recovered and annotated with two copies in the LrT2T genome, supported by both genomic and Iso-seq data. (**C**) The LrT2T genome corrects an assembly error involving two chromosomes in the previous version. The centromeric region of chromosome 3 (chr3) was previously unassembled due to densely clustered microsatellite sequences, resulting in its long and short arms being misassigned to two separate chromosomes (chr24 and chr64). The high coverage of both PacBio HiFi and ONT ultra-long reads enabled successful reconstruction of this centromeric region of chr3 in the LrT2T genome.

**Figure S5.**
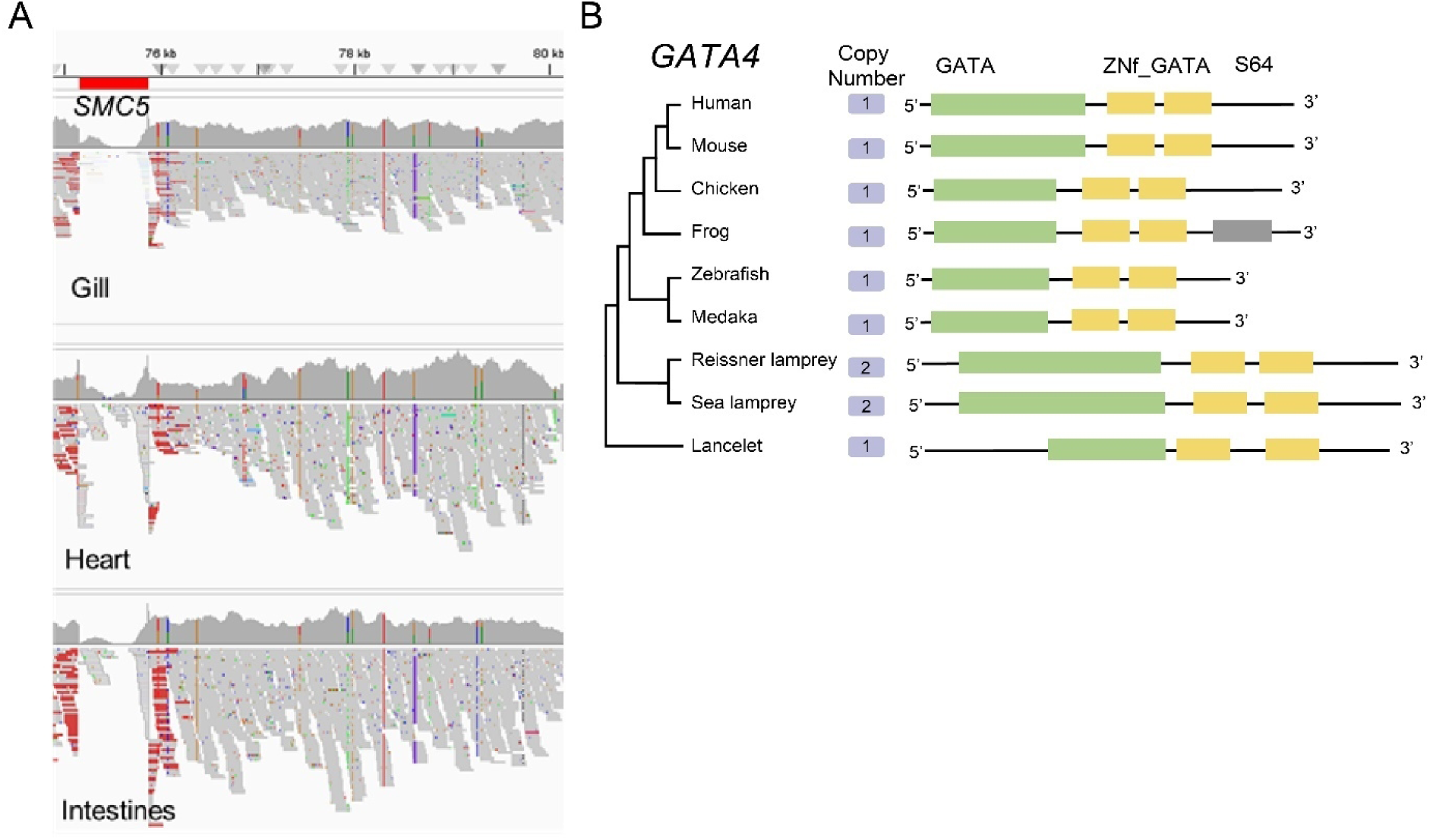
Loss and expansion of 2,797 newly annotated genes in LrT2T genome. (**A**) The *SMC5* gene region was lost in the genome of the lamprey. The aligned sequencing reads in the region (the red area) containing 10 *SMC5* copies on Chr1. There are gaps spanning the *SMC5* region in multiple somatic tissues, including gill, heart and intestines. (**B**) Evolutionary expansion of the *GATA4* gene family in lampreys (≥2 copies), in contrast to the single copy in other vertebrates. The conserved functional domains of *GATA4* in vertebrates are illustrated, including GATA-type transcription activator domain (GATA), Zinc finger DNA binding domain GATA-related (ZNf_GATA), and peptidase family S64 domains (S64).

**Figure S6.**
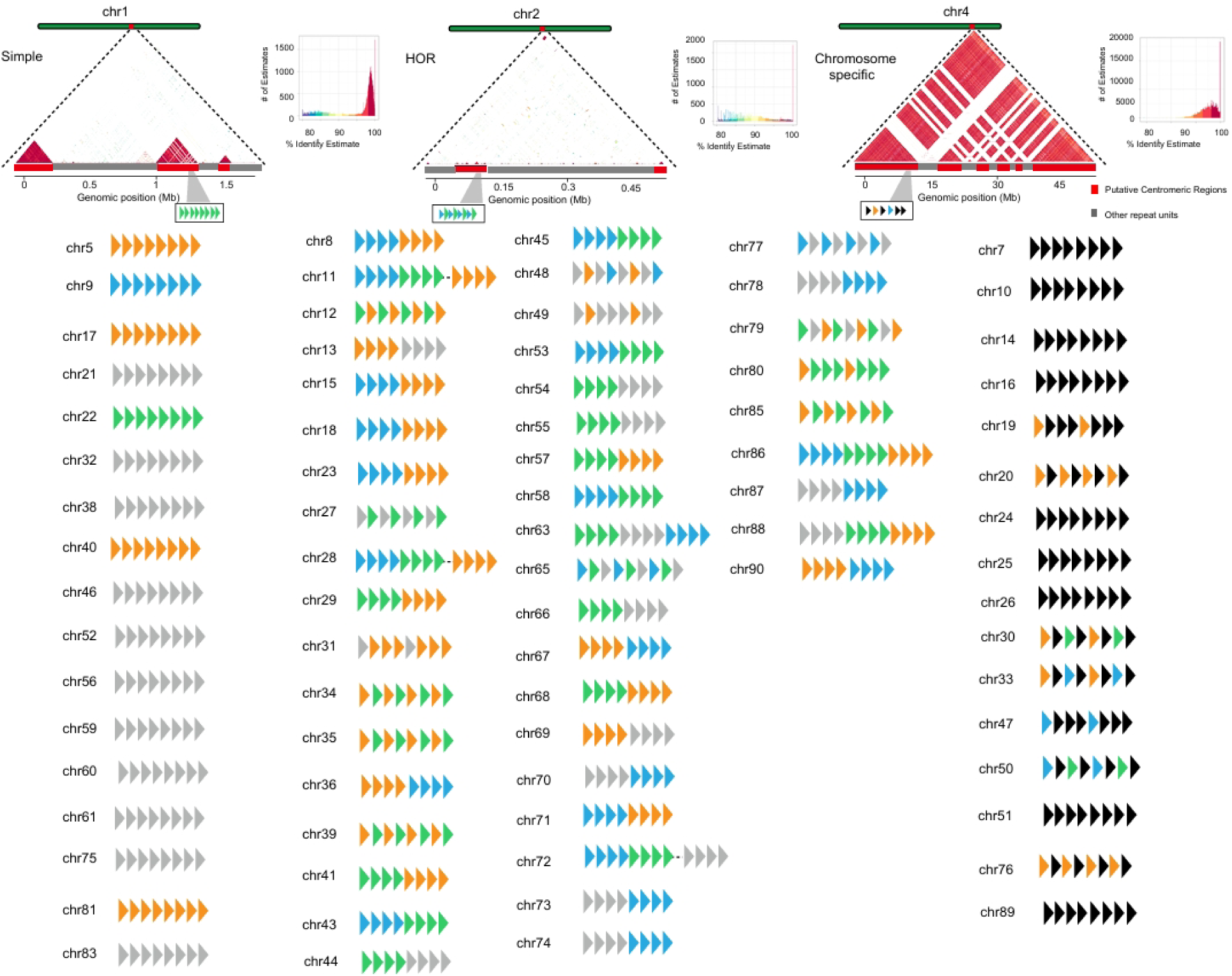
Centromere sequence The chromosome of lampry. Visualization and similarity scoring of representative centromeric regions from three chromosomes: the simple centromere chromosome 1, the HOR centromere chromosome 2, and chromosome specific centromere chromosome 4. The sequences of the tandem repeat units defining the centromeric regions are shown. The arrangement patterns of Lsat of the centromere sequences of other chromosomes, based on the three types of centromere sequences (simple, HOR and chromosome-specific), are presented.

**Figure S7.**
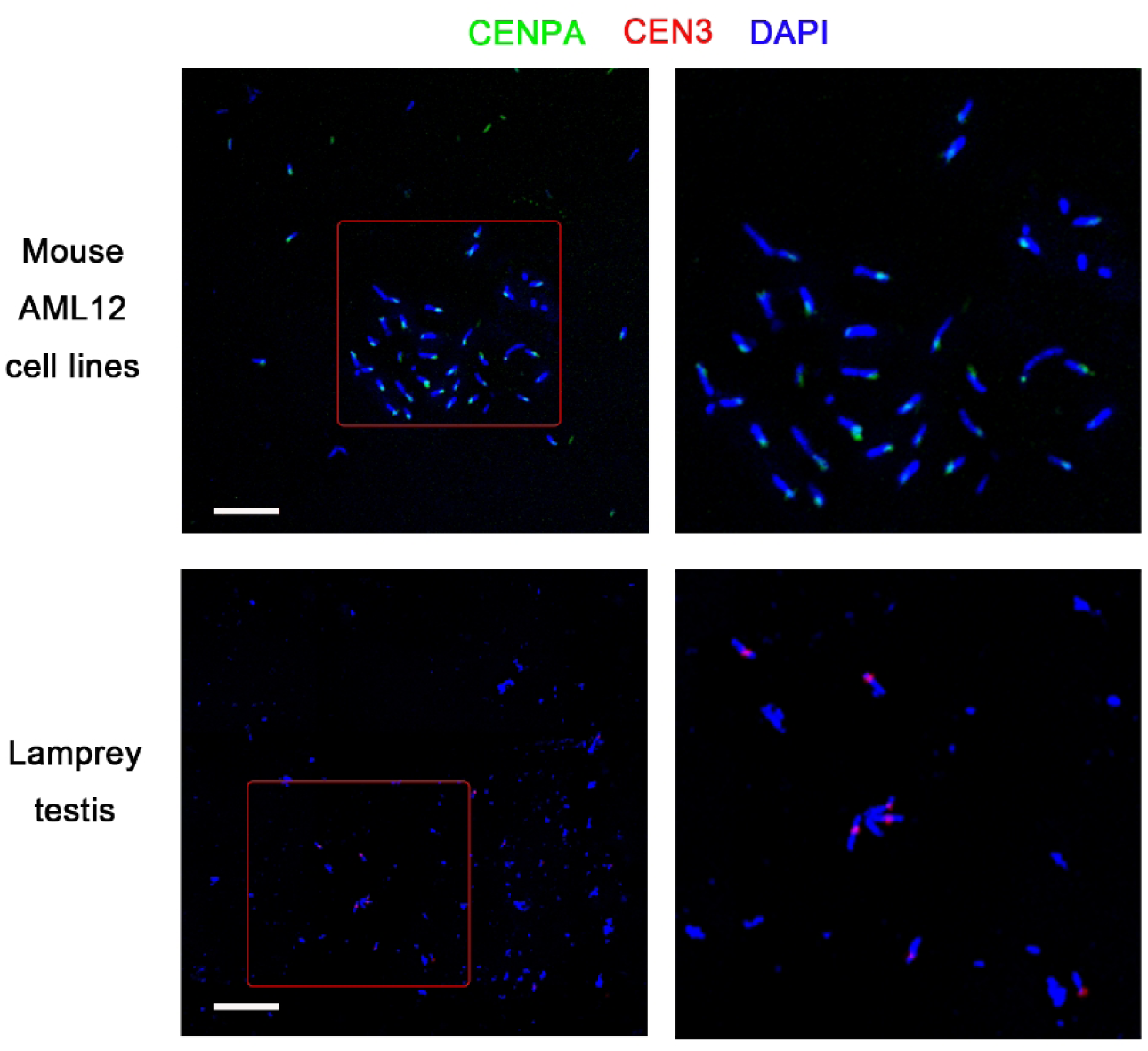
The chromosomal locations of CENPA and Cen-3. Mouse (AML12 lines, alpha mouse liver 1) and lamprey (primary cells from testis tissue) cells were used for immunofluorescence studies of CENPA and Cen3 to determine their localization information on chromosomes. The CENPA antibody is derived from abcam (EPR26697-213). The scale is 10 μm.

**Figure S8.**
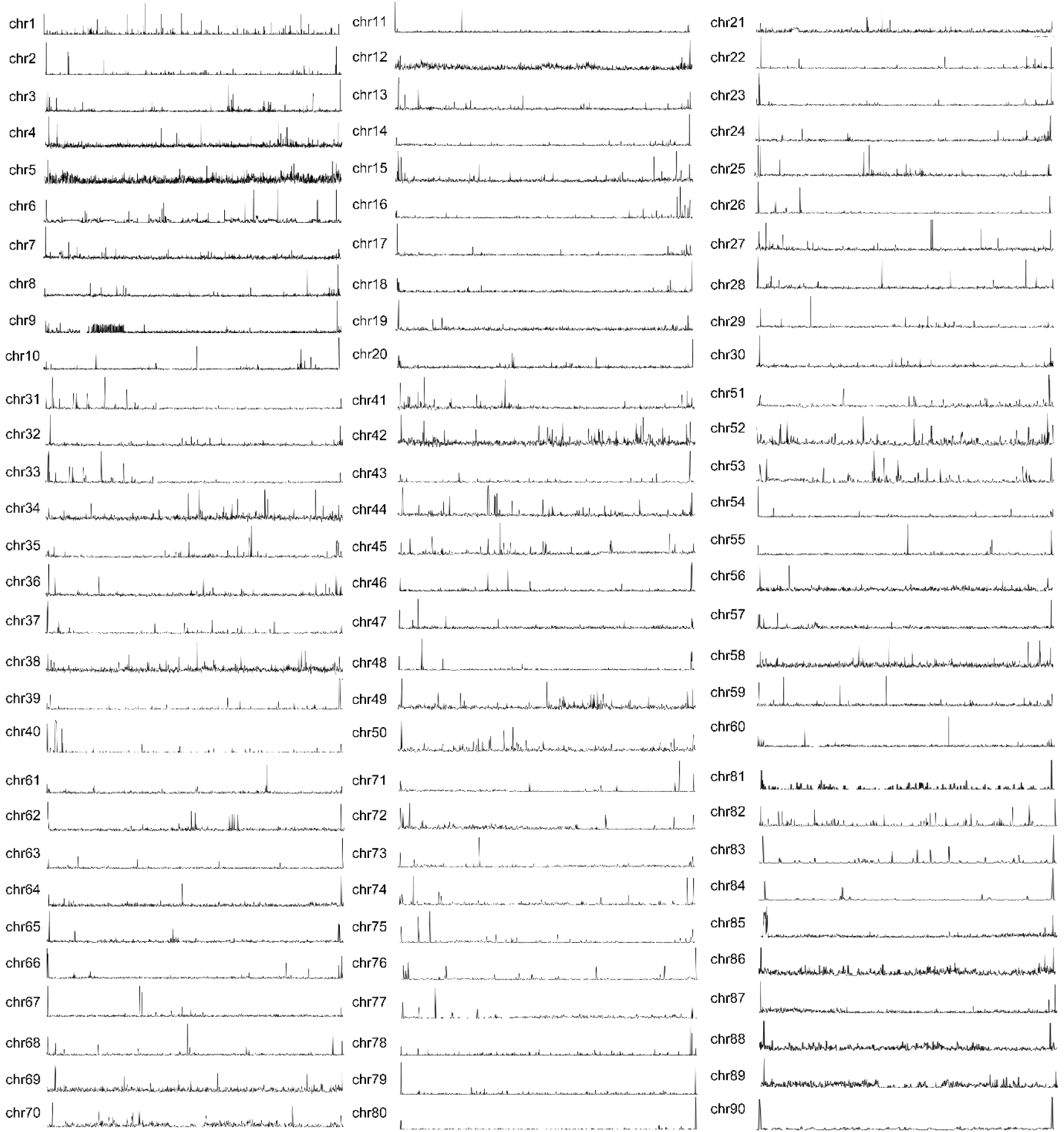
The distribution of telomere repeat sequences on the 90 chromosomes of the LrT2T genome. The height of the peak represents the number of telomere repeats. The conserved telomere sequence TTAGGG of vertebrates is assembled into each chromosome in the LrT2T genome.

**Figure S9.**
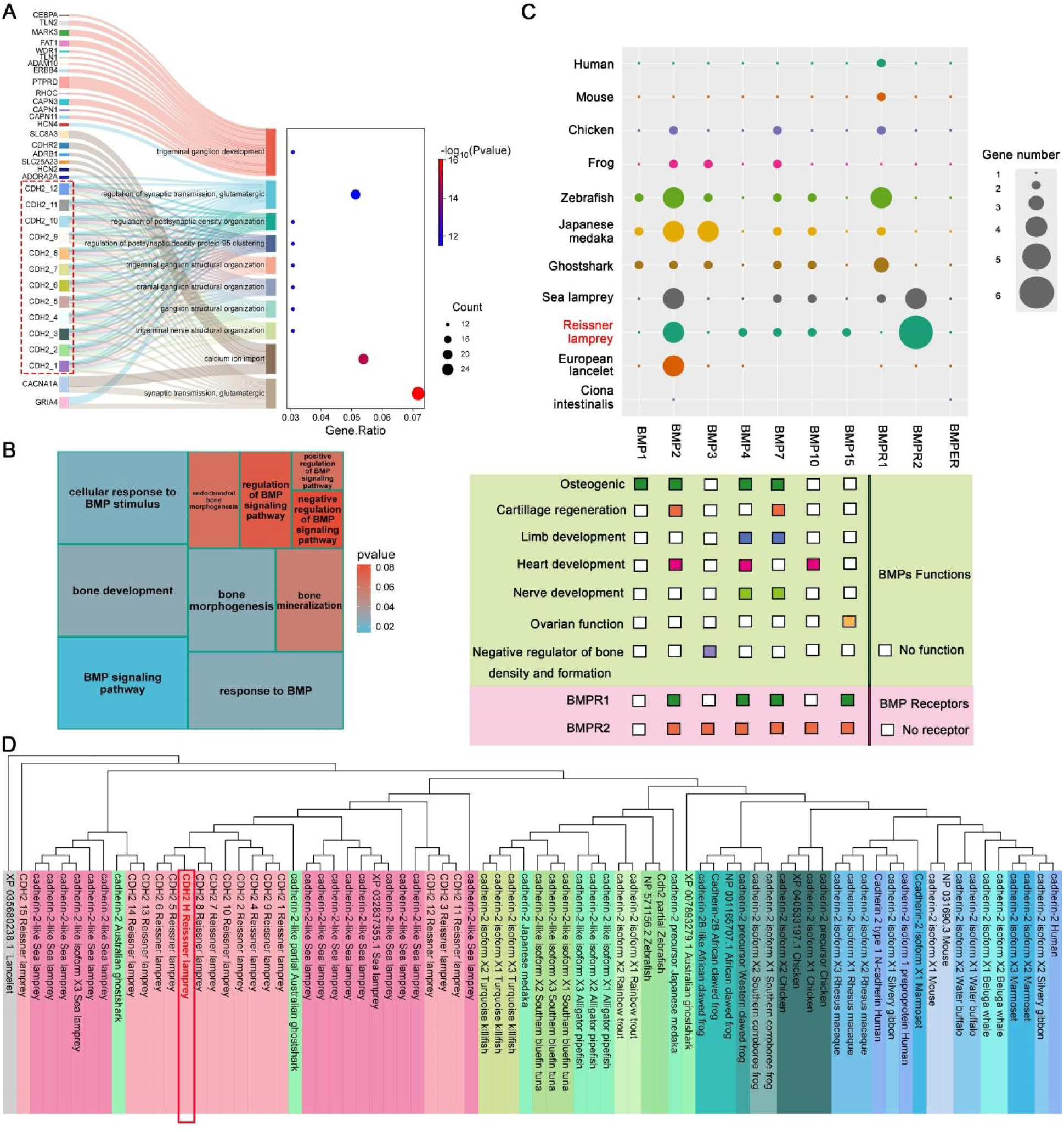
Gene family expansions associated with neural development and skeleton formation. (**A**) Enriched neurodevelopment-related GO terms identified from gene families expanded in the vertebrate common ancestor. (**B**) GO term enrichment for bone formation associated gene families expanded in the vertebrate common ancestor. (**C**) Copy number variation, functional categories, and receptor diversity of BMP signaling pathway components across vertebrate lineages. (**D**) Phylogenetic tree of *CDH2* gene copies across representative vertebrates, with lancelet as the outgroup. The tree was constructed using the maximum-likelihood method implemented in MEGA (12.0.11).

**Figure S10.**
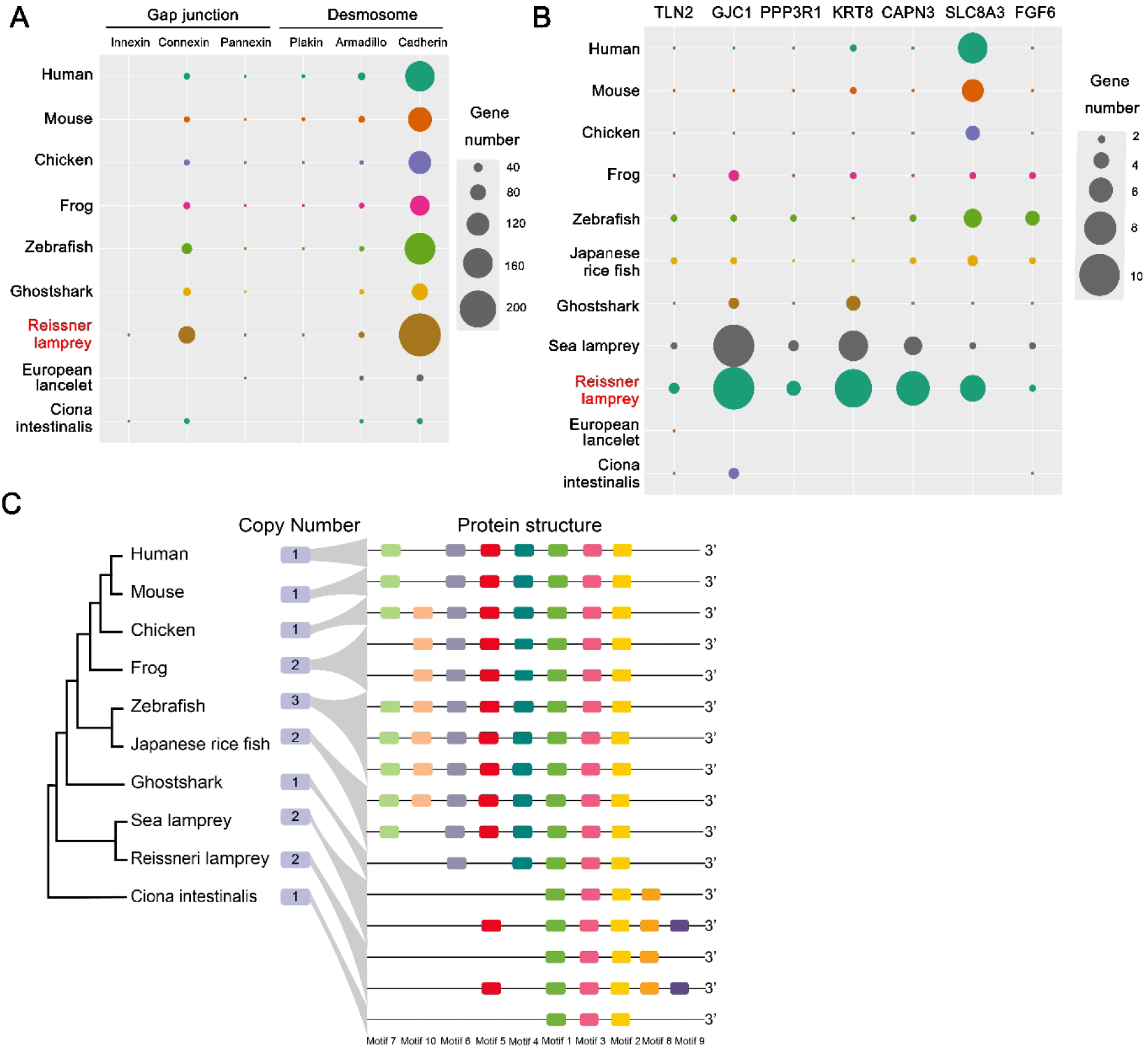
Expansions of gene families involved cell junction and cardiac function. (**A**) Expansion of the cell junction related gene family in vertebrates, including genes involved in gap junction and desmosome. (**B**) Expansion of other genes associated with cardiac function. (**C**) Evolution of gene copy number and protein functional motifs (Motif 1-9) of the *FGF6* gene family in vertebrates with ciona intestinalis as outgroup.

**Figure S11.**
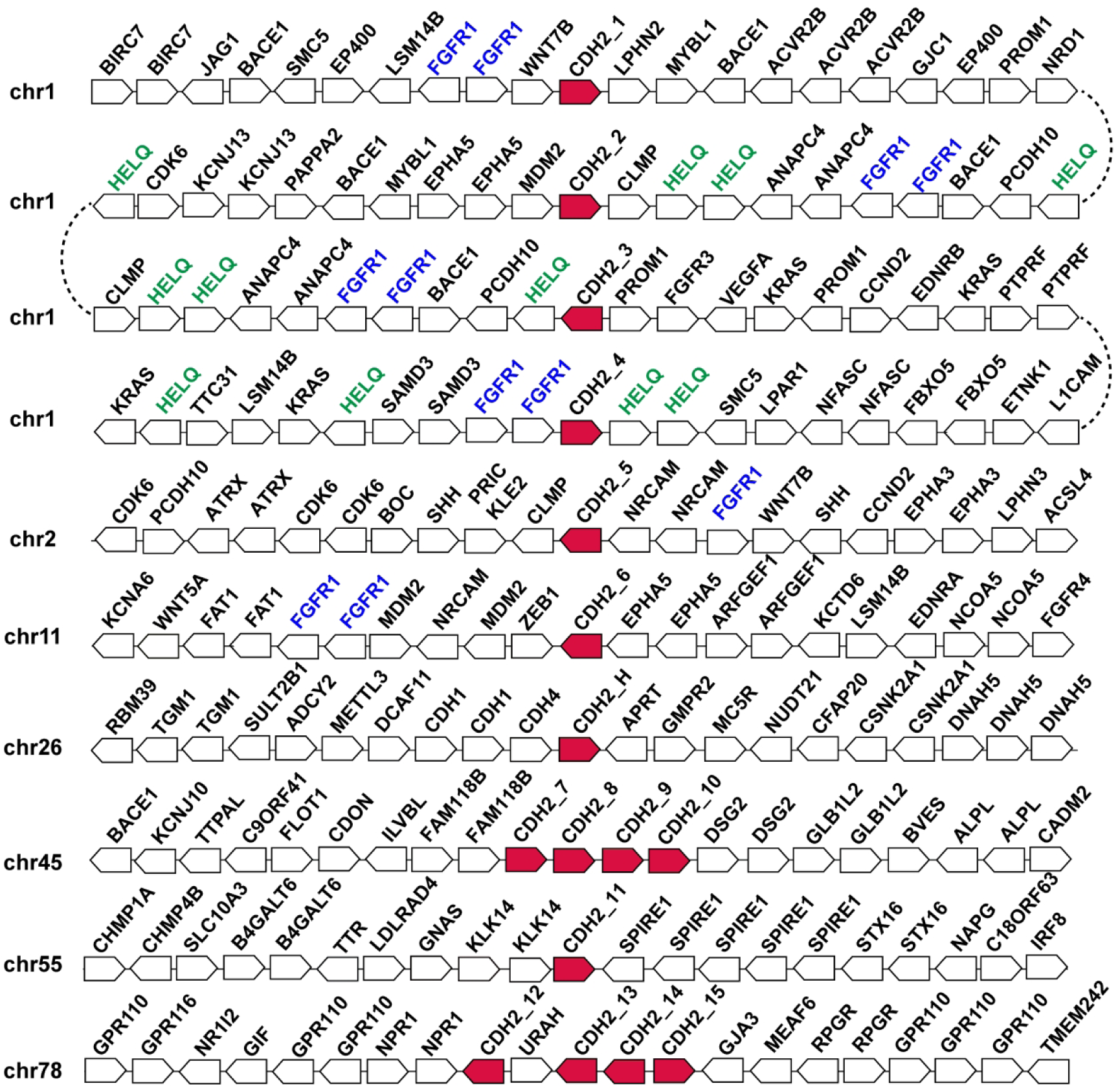
The distribution of all 16 copies of CDH2 in the LrT2T genome assembly. In Chr1 and Chr2, several genes (*HELQ* and *FGFR1*) appear frequently in the surroundings of *CDH2* copies, but the linkage patterns of other genes do not show clear sign of segmental duplications, implying a complex history of *CDH2* duplications that may involve multiple ancient structural changes.

**Figure S12.**
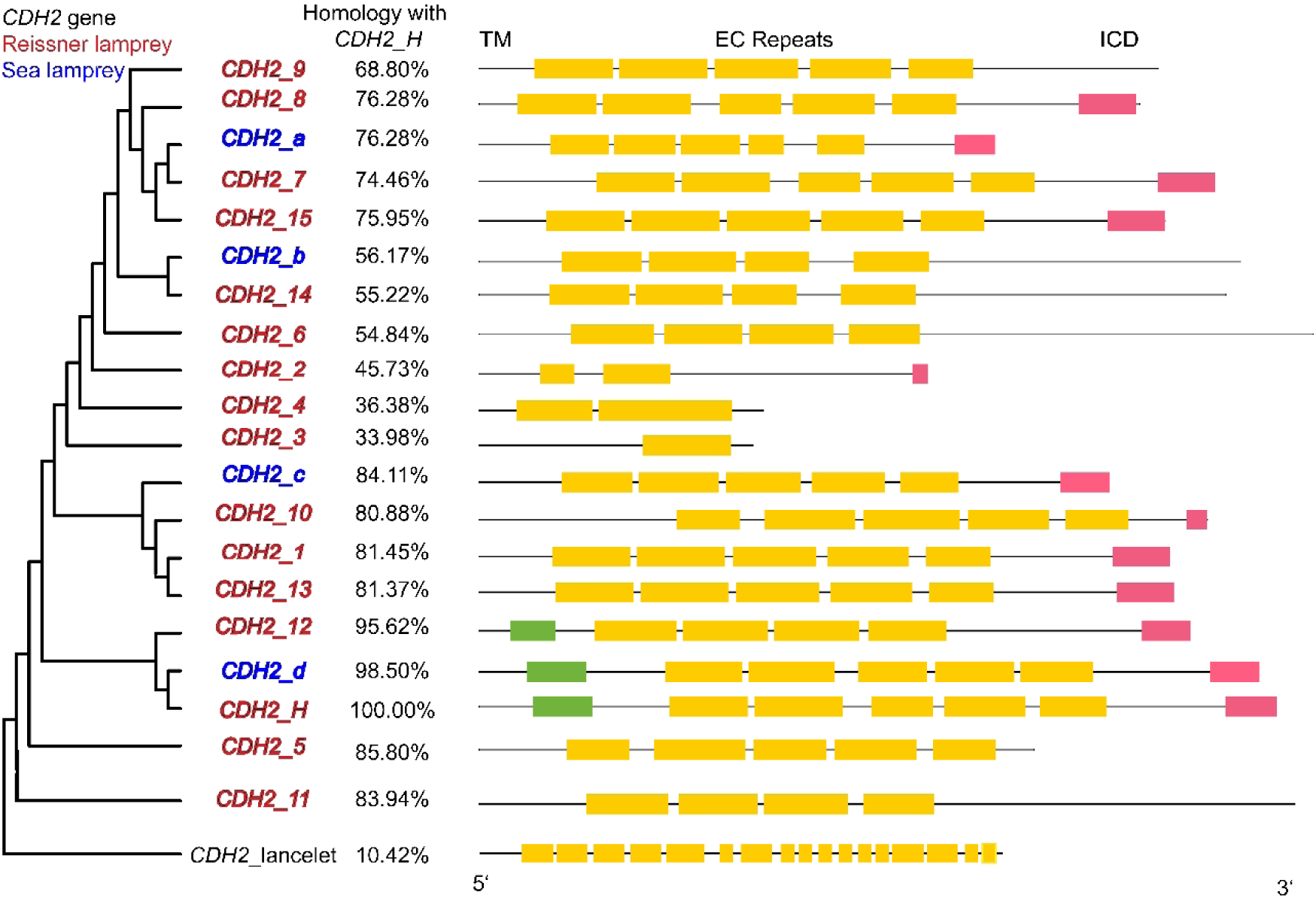
Phylogenetic tree of all copies of *CDH2* in LrT2T and sea lamprey. The sequence similarity among the 16 *CDH2* copies of *L. reissneri* ranges from 33.98%-98.50%. Among them, the *CDH2* sequence of the sea lamprey was selected from PMID: 40504158. Four copies were named *CDH2_a/b/c/d*. The conserved homology and functional domains of *CDH2* in lamprey are illustrated, including transmembrane domain (TM, green), extracellular cadherin (EC, yellow) repeats, and intracellular signaling domains (ICD, red). *CDH2_H* is close to the root node of the *CDH2* gene family, suggesting that *CDH2_H* is likely an early-emerging copy caused by the 1R event in the vertebrate common ancestor.

**Figure S13.**
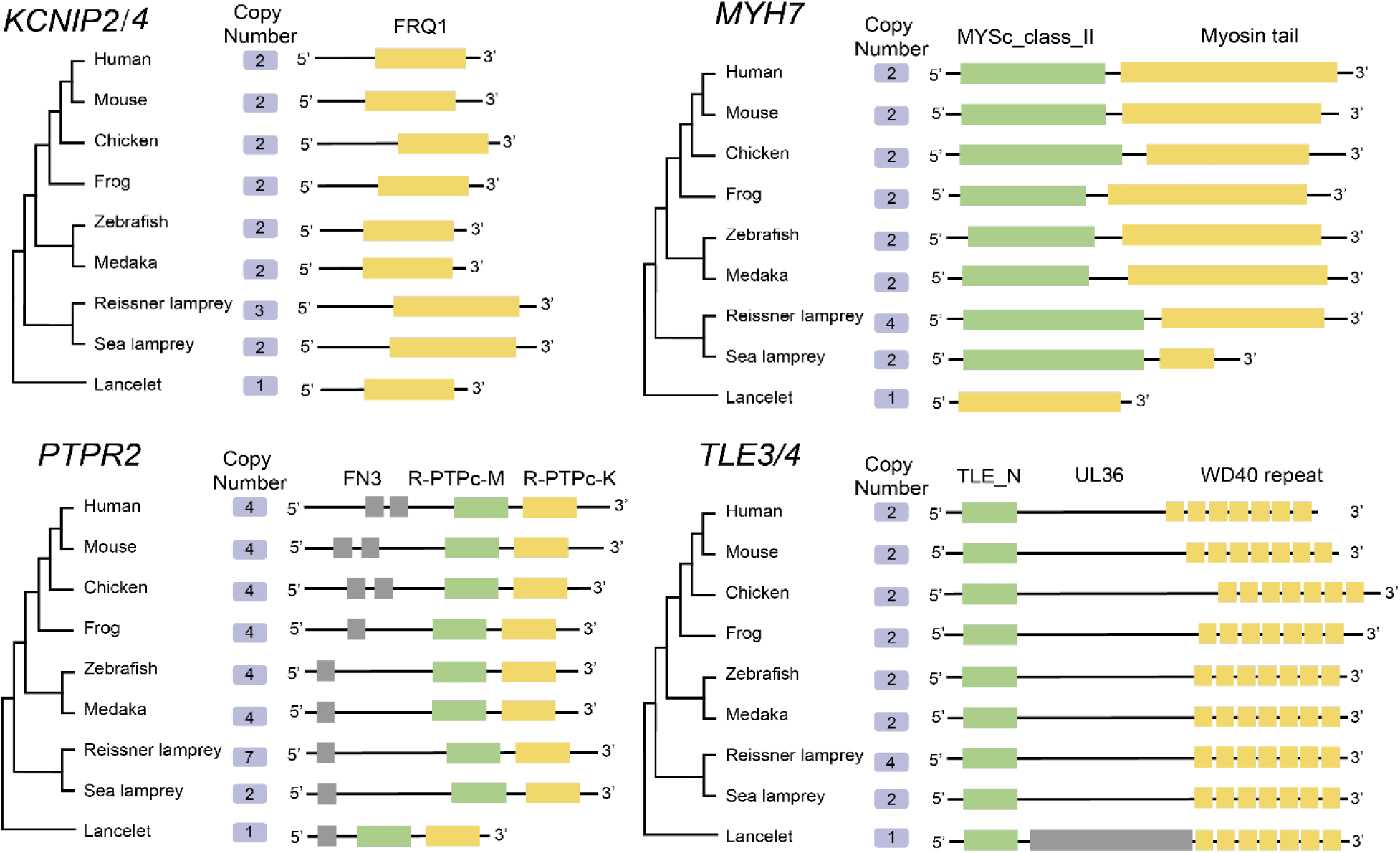
Evolution of the expansion genes identified in 1R in vertebrates with lancelet as outgroup. The gene copy number and conserved functional domains of 1,562 genes were identified in 1R in vertebrates are illustrated, including: EF-hand superfamily (FRQ1), class II myosins motor domain (MYSc_class_II), Myosin Tail Domain (Myosin Tail), catalytic domain of receptor-type tyrosine-protein phosphataseM (R-PTPc-M), PTP domain of receptor-type tyrosine-protein phosphatase K (R-PTPc-K), Fibronectin type III domain (FN3), Groucho/TLE N-terminal Q-rich domain (TLE_N), large tegument protein UL36 (UL36) and Trp-Asp (W-D).

**Figure S14.**
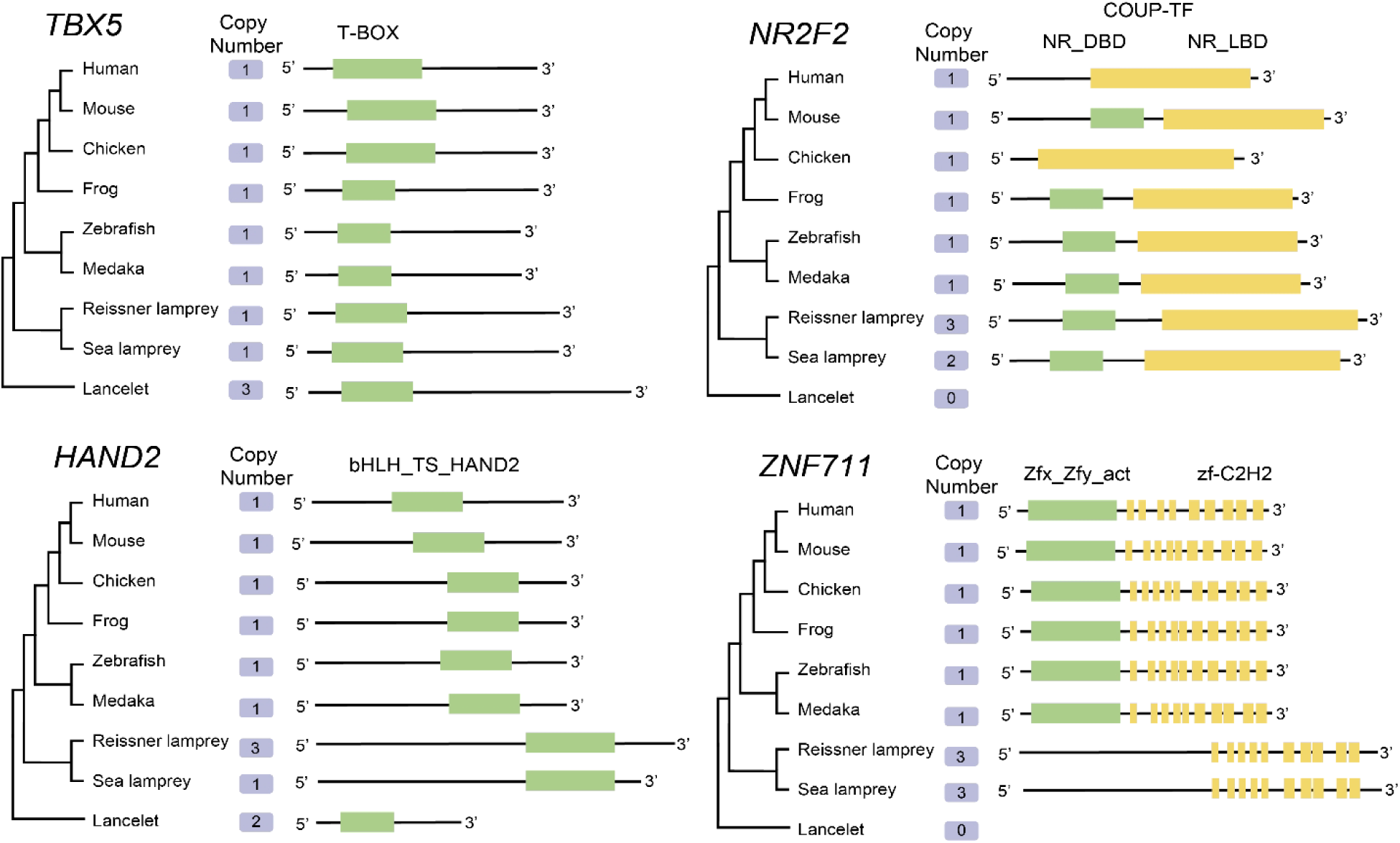
Evolution of the transcription factors related to cardiac chamber differentiation in vertebrates with lancelet as outgroup. The gene copy number and conserved functional domains of heart chamber formation related transcription factors in vertebrates are illustrated, including DNA-binding domain of T-box transcription factor (T-box), basic helix-loop-helix domain found in heart- and neural crest derivatives-expressed protein 2 and similar proteins (bHLH_TS_HAND2), ligand binding domain of chicken ovalbumin upstream promoter transcription factors (NR_LBD), DNA-binding domain of chicken ovalbumin upstream promoter transcription factors (NR_DBD), Zfx / Zfy transcription activation region (Zfx_Zfy_act) and Zinc finger, C2H2 type (zf-C2H2). The gene copy number indicates that *TBX5* did not undergo expansion, while *HAND2*, *NR2F2*, and *ZNF711* underwent a single doubling (CR) in the lamprey genome. In addition, *NR2F2* and *ZNF711* may be new genes formed in the genome of lampreys.

**Figure S15.**
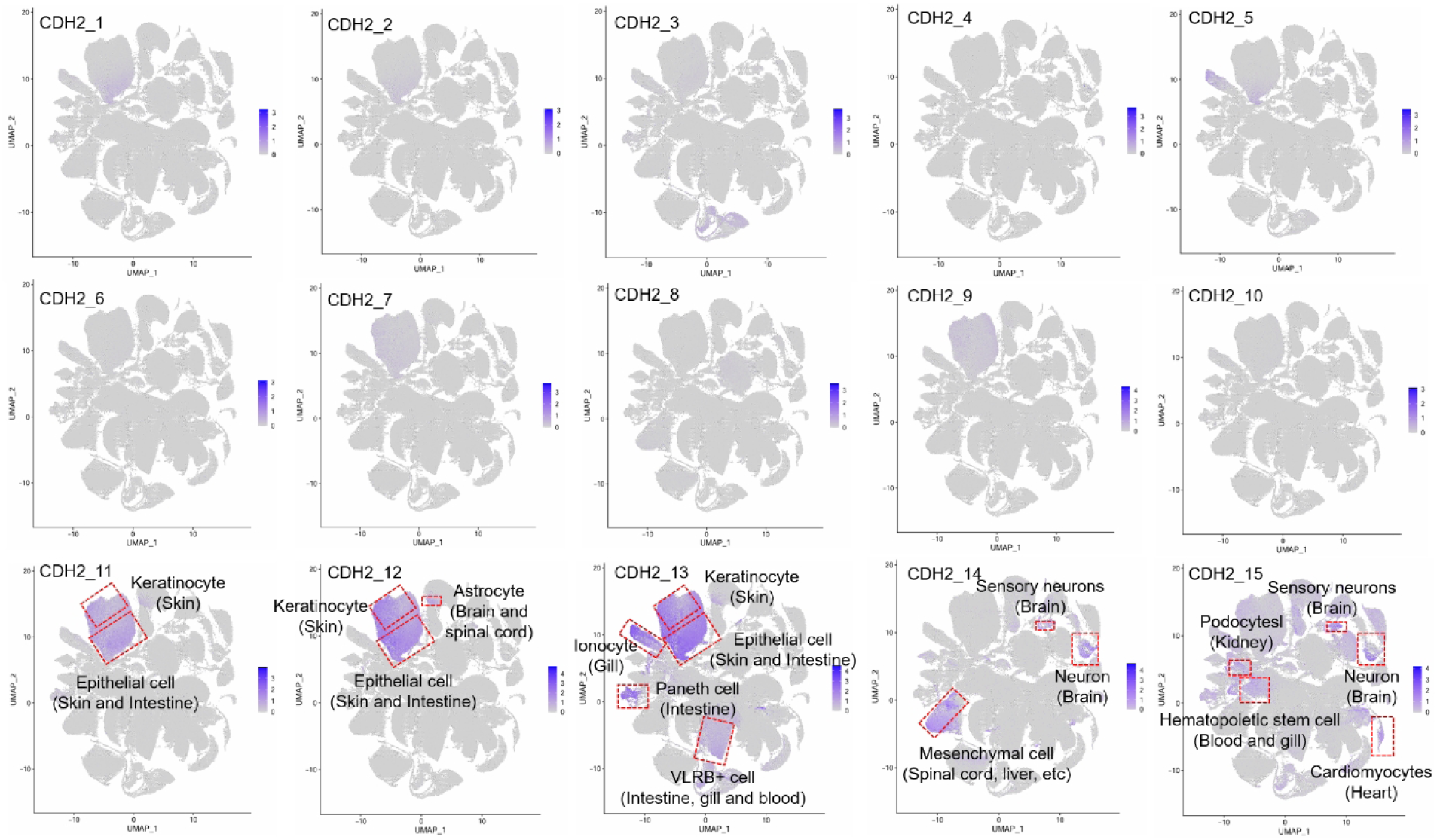
Single-cell atlas of lamprey showing expression patterns of 15 *CDH2* gene copies across diverse cell types. Cell types and their tissue origins are indicated for *CDH2* copies exhibiting cell type-associated expression patterns.

**Figure S16.**
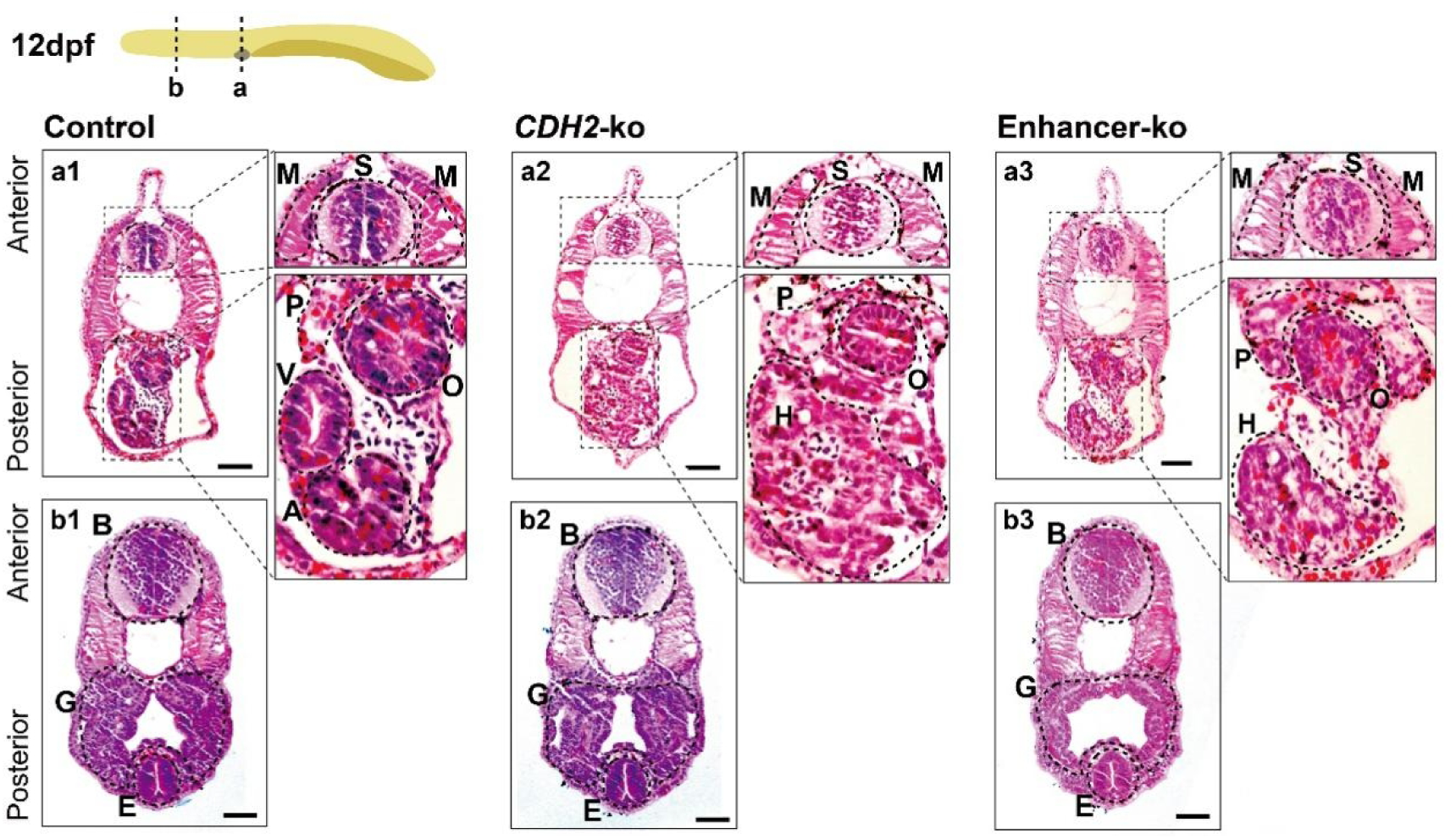
Histological assessment of heart chamber structure and cardiomyocyte organization by H&E staining at 12 dpf. The magnified area is a typical cardiac region (A-atrium, V-ventricle), spinal cord (S), muscle (M), pronephros (P), oesophagus (O), brain (B), gill (G) and endostyle (E), and the scale is 50 μm.

**Table S1.** Statistics of the sequencing reads used for lamprey genome assembly.

| Term | Total bases (Gb) | Sequencing depth (×) | N50 (kb) |
| --- | --- | --- | --- |
| PacBio HiFi | 114.3 | 91.4 | 16.5 |
| Ultra-long ONT | 139.2 | 111.4 | 100.2 |
| Hi-C | 254.9 | 101.9 |  |
| Illumina | 125.7 | 100.6 |  |

Note: The statistics was generated using the filtered reads. The sequencing depth was calculated based on the assembled genome size.

**Table S2.**
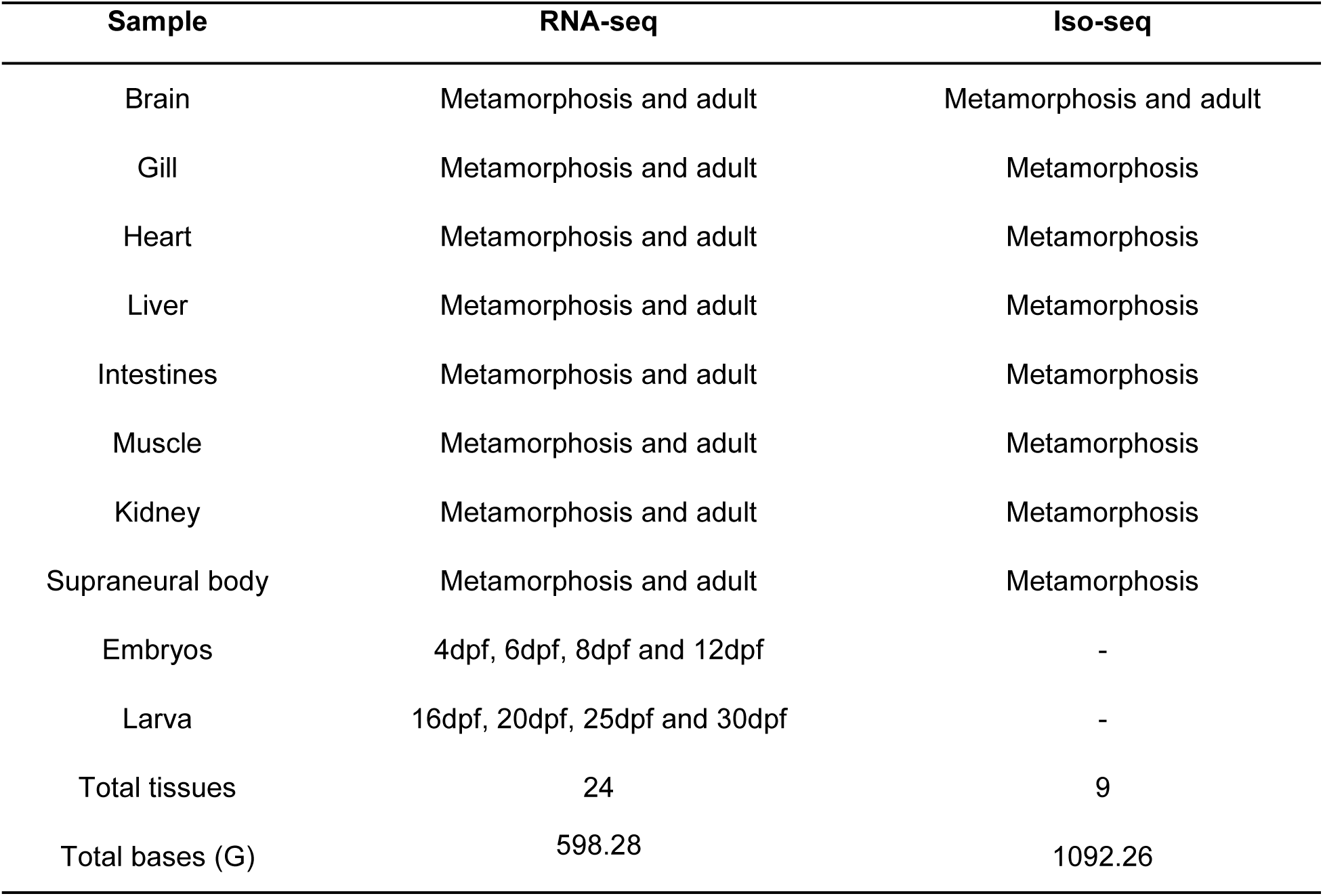
Sampled organs from different developmental stages for RNA-seq and Iso-seq.

| Sample | RNA-seq | Iso-seq |
| --- | --- | --- |
| Brain | Metamorphosis and adult | Metamorphosis and adult |
| Gill | Metamorphosis and adult | Metamorphosis |
| Heart | Metamorphosis and adult | Metamorphosis |
| Liver | Metamorphosis and adult | Metamorphosis |
| Intestines | Metamorphosis and adult | Metamorphosis |
| Muscle | Metamorphosis and adult | Metamorphosis |
| Kidney | Metamorphosis and adult | Metamorphosis |
| Supraneural body | Metamorphosis and adult | Metamorphosis |
| Embryos | 4dpf, 6dpf, 8dpf and 12dpf | - |
| Larva | 16dpf, 20dpf, 25dpf and 30dpf | - |
| Total tissues | 24 | 9 |
| Total bases (G) | 598.28 | 1092.26 |

**Table S3.**
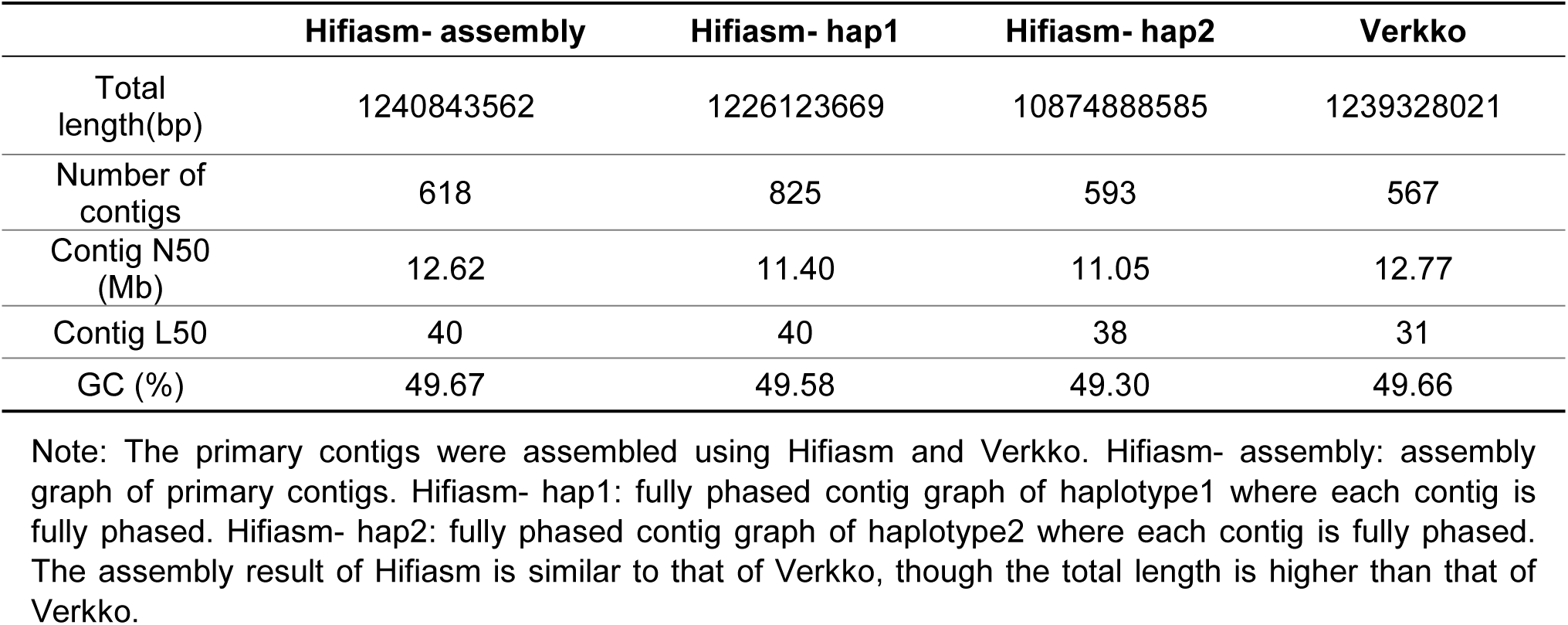
The quality of the draft genome after the primary contigs by Hifiasm and Verkko.

|  | Hifiasm- assembly | Hifiasm- hap1 | Hifiasm- hap2 | Verkko |
| --- | --- | --- | --- | --- |
| Total length(bp) | 1240843562 | 1226123669 | 10874888585 | 1239328021 |
| Number of contigs | 618 | 825 | 593 | 567 |
| Contig N50 (Mb) | 12.62 | 11.40 | 11.05 | 12.77 |
| Contig L50 | 40 | 40 | 38 | 31 |
| GC (%) | 49.67 | 49.58 | 49.30 | 49.66 |

Note: The primary contigs were assembled using Hifiasm and Verkko. Hifiasm- assembly: assembly graph of primary contigs. Hifiasm- hap1: fully phased contig graph of haplotype1 where each contig is fully phased. Hifiasm- hap2: fully phased contig graph of haplotype2 where each contig is fully phased. The assembly result of Hifiasm is similar to that of Verkko, though the total length is higher than that of Verkko.

**Table S4.** Basic statistics of different genome (reissner lamprey) assembly versions.

|  | Primary assembly | Hi-C scaffolds | Complete assembly |
| --- | --- | --- | --- |
| Total number ($\geq 0$ Bp) | 618 | 182 | 182 |
| Total number ($\geq 1$ Kb) | 595 | 182 | 182 |
| Total number ($\geq 5$ Kb) | 567 | 180 | 180 |
| Total number ($\geq 10$ Kb) | 557 | 179 | 179 |
| Total number ($\geq 25$ Kb) | 530 | 170 | 170 |
| Total number ($\geq 50$ Kb) | 4470 | 157 | 157 |
| Total length ($\geq 0$ Bp) | 1,240,843,562 | 1,240,917,391 | 1,255,687,010 |
| Total length ($\geq 1$ Kb) | 1,240,833,508 | 1,240,917,391 | 1,255,687,010 |
| Total length ($\geq 5$ Kb) | 1,240,756,330 | 1,240,912,704 | 1,255,682,323 |
| Total length ($\geq 10$ Kb) | 1,240,678,898 | 1,240,903,010 | 1,255,672,629 |
| Total length ($\geq 25$ Kb) | 1,240,136,585 | 1,240,728,438 | 1,255,498,067 |
| Total length ($\geq 50$ Kb) | 1,237,876,005 | 1,240,272,238 | 1,255,043,823 |
| GC (%) | 49.67 | 49.66 | 49.68 |
| N50 | 12,624,862 | 13,411,000 | 13,411,051 |
| N90 | 2,012,395 | 8,178,101 | 8,178,080 |
| L50 | 40 | 31 | 31 |
| L90 | 120 | 75 | 75 |

Note: N50 – The length of the shortest scaffold such that the combined length of all scaffolds of this size or longer constitutes at least 50 % of the total genome assembly. N90 – The length of the shortest scaffold such that the combined length of all scaffolds of this size or longer constitutes at least 90 % of the total genome assembly. L50 – The minimum number of longest scaffolds required to reach 50 % of the total assembly length. L90 – The minimum number of longest scaffolds required to reach 90 % of the total assembly length.

**Table S5.** Statistics of the assembled chromosome-level genome using Hi-C data.

| Chromosome ID | Length (bp) | Chromosome ID | Length (bp) | Chromosome ID | Length (bp) |
| --- | --- | --- | --- | --- | --- |
| HiC_chr_1 | 92,830,426 | HiC_chr_31 | 13,411,051 | HiC_chr_61 | 10,600,573 |
| HiC_chr_2 | 38,771,648 | HiC_chr_32 | 13,379,768 | HiC_chr_62 | 10,262,188 |
| HiC_chr_3 | 33,179,969 | HiC_chr_33 | 13,376,271 | HiC_chr_63 | 10,148,103 |
| HiC_chr_4 | 25,474,541 | HiC_chr_34 | 13,351,348 | HiC_chr_64 | 9,823,262 |
| HiC_chr_5 | 24,706,506 | HiC_chr_35 | 13,314,242 | HiC_chr_65 | 9,786,390 |
| HiC_chr_6 | 25,651,993 | HiC_chr_36 | 13,311,383 | HiC_chr_66 | 9,658,798 |
| HiC_chr_7 | 23,043,219 | HiC_chr_37 | 13,188,733 | HiC_chr_67 | 9,258,961 |
| HiC_chr_8 | 21,462,600 | HiC_chr_38 | 13,141,897 | HiC_chr_68 | 9,234,072 |
| HiC_chr_9 | 20,867,936 | HiC_chr_39 | 13,139,516 | HiC_chr_69 | 9,209,501 |
| HiC_chr_10 | 18,562,780 | HiC_chr_40 | 13,073,196 | HiC_chr_70 | 8,772,014 |
| HiC_chr_11 | 19,392,343 | HiC_chr_41 | 13,029,938 | HiC_chr_71 | 8,477,365 |
| HiC_chr_12 | 16,359,990 | HiC_chr_42 | 12,972,513 | HiC_chr_72 | 8,469,877 |
| HiC_chr_13 | 16,242,513 | HiC_chr_43 | 12,866,419 | HiC_chr_73 | 8,431,400 |
| HiC_chr_14 | 15,977,733 | HiC_chr_44 | 12,850,073 | HiC_chr_74 | 8,289,389 |
| HiC_chr_15 | 15,802,445 | HiC_chr_45 | 12,846,757 | HiC_chr_75 | 8,178,080 |
| HiC_chr_16 | 15,793,521 | HiC_chr_46 | 12,601,181 | HiC_chr_76 | 7,882,660 |
| HiC_chr_17 | 15,793,074 | HiC_chr_47 | 12,450,408 | HiC_chr_77 | 7,847,768 |
| HiC_chr_18 | 15,671,823 | HiC_chr_48 | 12,006,100 | HiC_chr_78 | 7,783,700 |
| HiC_chr_19 | 15,578,989 | HiC_chr_49 | 11,993,630 | HiC_chr_79 | 7,613,853 |
| HiC_chr_20 | 15,022,049 | HiC_chr_50 | 11,522,324 | HiC_chr_80 | 7,741,159 |
| HiC_chr_21 | 14,939,785 | HiC_chr_51 | 11,474,880 | HiC_chr_81 | 7,522,394 |
| HiC_chr_22 | 14,516,591 | HiC_chr_52 | 11,213,828 | HiC_chr_82 | 7,448,177 |
| HiC_chr_23 | 14,461,270 | HiC_chr_53 | 11,169,690 | HiC_chr_83 | 7,381,794 |
| HiC_chr_24 | 14,402,432 | HiC_chr_54 | 11,000,975 | HiC_chr_84 | 7,318,088 |
| HiC_chr_25 | 14,283,824 | HiC_chr_55 | 10,987,781 | HiC_chr_85 | 7,299,022 |
| HiC_chr_26 | 13,994,013 | HiC_chr_56 | 10,824,243 | HiC_chr_86 | 7,103,495 |
| HiC_chr_27 | 13,889,567 | HiC_chr_57 | 10,819,341 | HiC_chr_87 | 6,999,759 |
| HiC_chr_28 | 13,798,269 | HiC_chr_58 | 10,782,189 | HiC_chr_88 | 6,755,461 |
| HiC_chr_29 | 13,680,316 | HiC_chr_59 | 10,669,597 | HiC_chr_89 | 6,090,654 |
| HiC_chr_30 | 13,582,357 | HiC_chr_60 | 10,654,661 | HiC_chr_90 | 3,448,655 |
| Total chromosome level |  |  |  | 1,239,995,067 |  |
| Total length |  |  |  | 1,255,687,010 |  |
| Chromosome/total |  |  |  | 98.75 % |  |

**Table S6.**
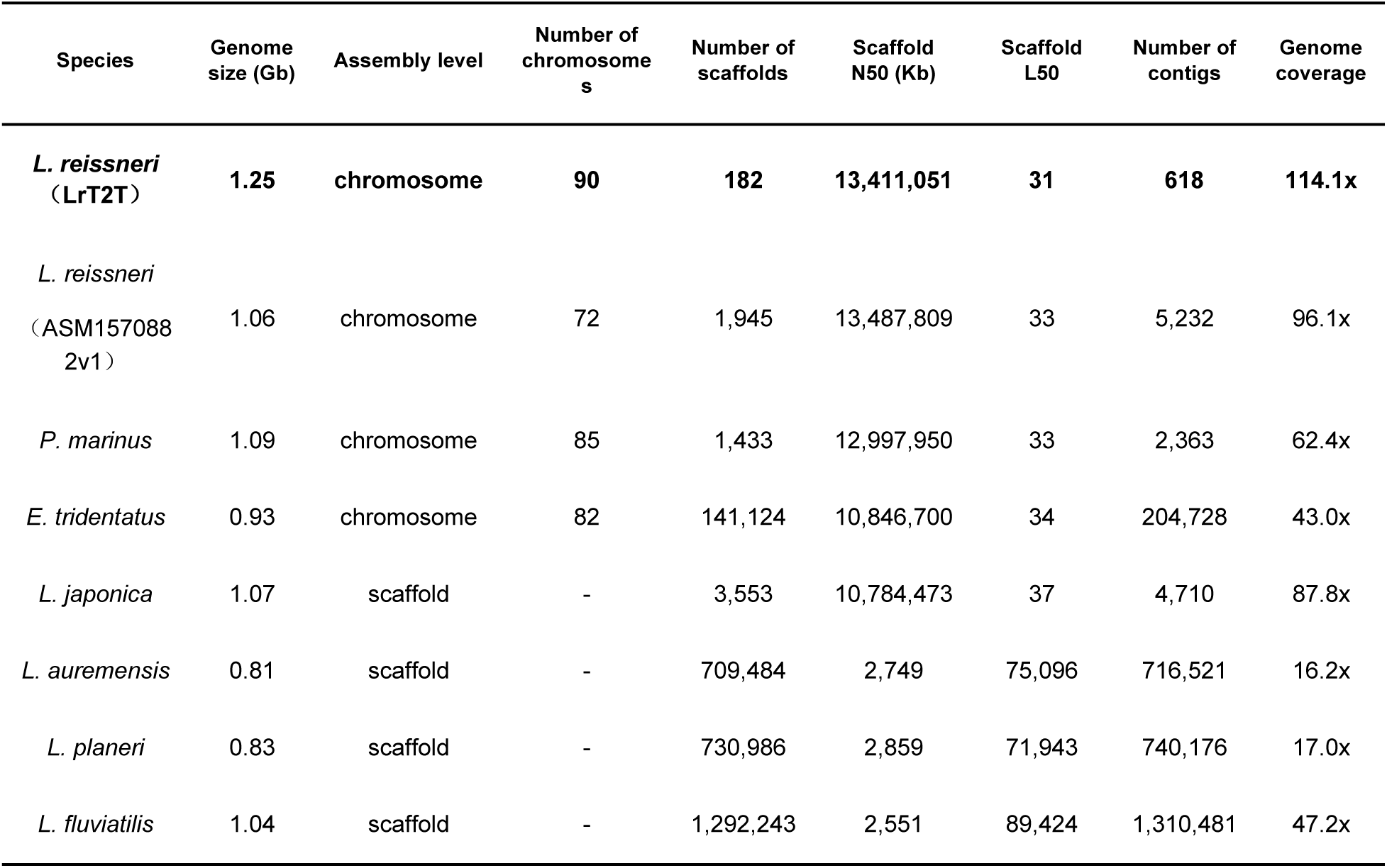
Comparison of LrT2T with the released lamprey genomes.

**Table S7.** BUSCO results of the lamprey genome.

| Library | Complete BUSCOs (C) | Complete and single-copy BUSCOs (S) | Complete and duplicated BUSCOs (D) | Fragmented BUSCOs (F) | Missing BUSCOs (M) | Total BUSCO groups searched | Complete BUSCOs (%) |
| --- | --- | --- | --- | --- | --- | --- | --- |
| *L. reissneri* (LrT2T) | 904 | 879 | 25 | 29 | 21 | 954 | 94.7 |
| *L. reissneri* (ASM1570882v1) | 788 | 720 | 68 | 30 | 136 | 954 | 82.6 |
| *P. marinus* | 882 | 863 | 19 | 41 | 31 | 954 | 92.5 |
| *E. tridentatus* | 369 | 364 | 5 | 353 | 232 | 954 | 38.7 |
| *L. japonica* | 810 | 745 | 65 | 44 | 100 | 954 | 84.9 |
| *L. auremensis* | 320 | 319 | 1 | 286 | 248 | 954 | 33.5 |
| *L. planeri* | 349 | 347 | 2 | 351 | 254 | 954 | 36.6 |
| *L. fluviatilis* | 718 | 707 | 11 | 91 | 145 | 954 | 75.3 |

Note: The genome integrity of the publicly available lamprey genome was compared using BUSCO. The reference orthologous groups are from https://busco.ezlab.org/.

**Table S8.**
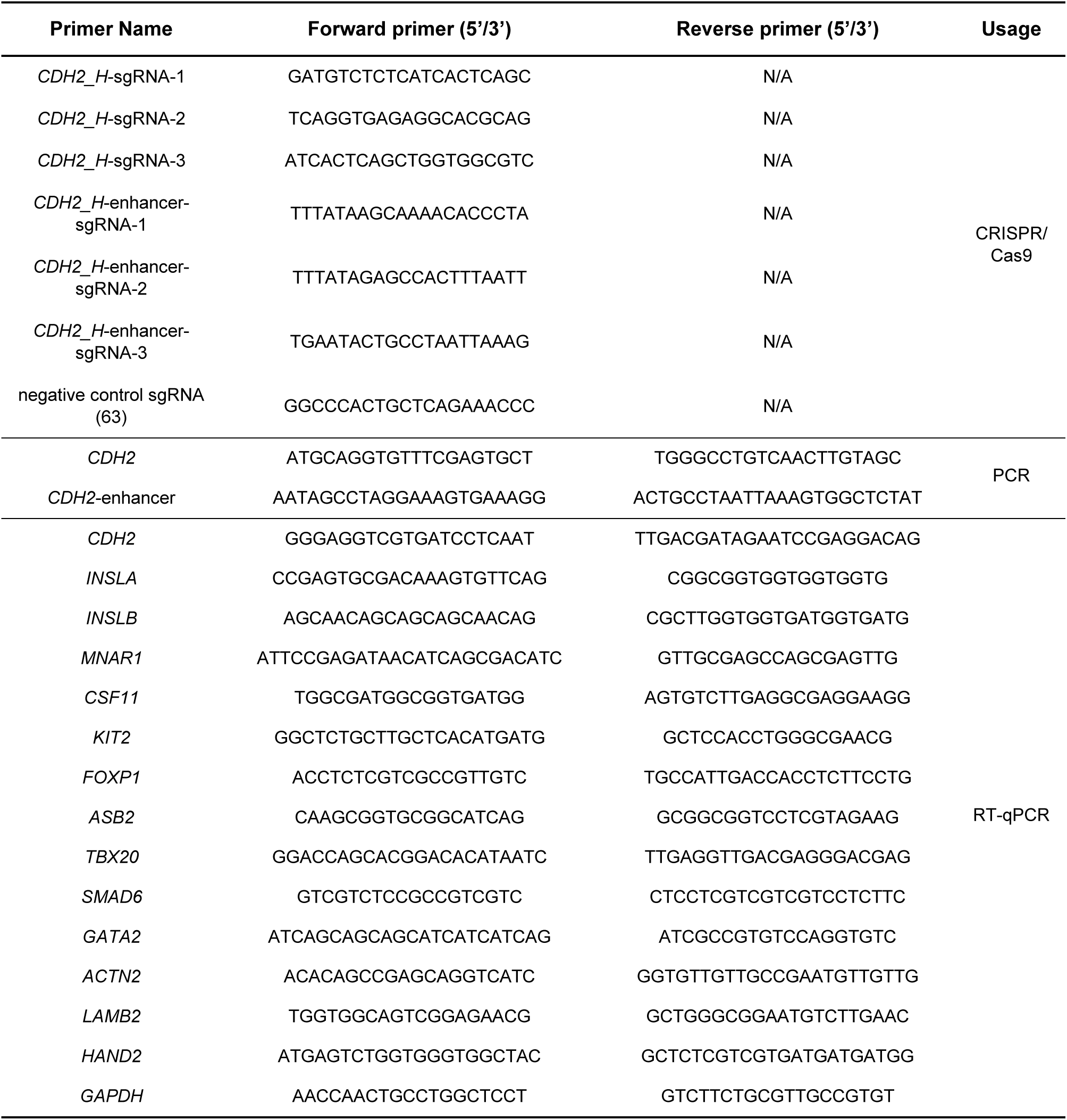
Sequence information of the primers used in this study.

## Notes

### Competing Interest Statement

The authors have declared no competing interest.

